# EchinoDB: An update to the web-based application for genomic and transcriptomic data on echinoderms

**DOI:** 10.1101/2022.01.03.474134

**Authors:** Varnika Mittal, Robert W. Reid, Denis Jacob Machado, Vladimir Mashanov, Daniel A. Janies

## Abstract

Here we release a new version of EchinoDB (https://echinodb.uncc.edu). EchinoDB is a database of genomic and transcriptomic data on echinoderms. The initial database consisted of groups of 749,397 orthologous and paralogous transcripts arranged in orthoclusters by sequence similarity. The new version of EchinoDB includes RNA-seq data of the brittle star *Ophioderma brevispinum* and high-quality genomic assembly data of the green sea urchin *Lytechinus variegatus*. In addition, we enabled keyword searches for annotated data and installed an updated version of Sequenceserver to allow BLAST searches. The data are downloadable in FASTA format. The first version of EchinoDB appeared in 2016 and was implemented in GO on a local server. The new version has been updated using R Shiny to include new features and improvements in the application. Furthermore, EchinoDB now runs entirely in the cloud for increased reliability and scaling. EchinoDB enjoys a user base drawn from the fields of phylogenetics, developmental biology, genomics, physiology, neurobiology, and regeneration. As use cases, we illustrate how EchinoDB is used in discovering pathways and gene regulatory networks involved in the tissue regeneration process.

## Introduction

EchinoDB (1) is an open-source web-based application accessed via https://echinodb.uncc.edu (19). An effective bioinformatics tool must keep up with new data, advances in software, server architecture, and programming languages. To scale well with increasing amounts of data, reliability and number of users, we have updated EchinoDB to take advantage of cloud services. We have also added new datasets along with new features such as sequence downloading functionality, Basic Local Alignment Search Tool (BLAST) (13) and keyword search to efficiently query the annotated data.

EchinoDB is a repository of genomic and amino acid sequence data from across the phylum Echinodermata. The primary dataset is organized into clusters of orthologous genes (orthoclusters). Additional datasets include RNA-seq data of the brittle star *Ophioderma brevispinum* (Say, 1825) (Echinodermata: Ophiuroidea: Ophiacanthida: Ophiodermatidae) (5) and genome assembly data of green sea urchin *Lytechinus variegatus* (Lamarck, 1816) (Echinodermata: Echinoidea: Camarodonta: Toxopneustidae) (6).

EchinoDB serves a large and diverse community of researchers focused on different aspects of biology including phylogenetics, developmental biology, genomics, physiology, neurobiology, and regeneration. As use cases, we illustrate how EchinoDB is used in discovering pathways and gene regulatory networks involved in the tissue regeneration process.

Echinoderms are a phylum of marine invertebrate deuterostomes and thus share a deep common ancestor with vertebrates (9, HYPERLINK \l "bookmark12" 10, HYPERLINK \l "bookmark16" 14). However, unlike vertebrates, many echinoderm species can regenerate all their tissue types after injury without inducing cancers (15). Therefore, with the correct resources, including genomic and transcriptomic data, echinoderms can be developed as a model organism to research regulatory mechanisms for regeneration.

The raw data in EchinoDB were generated by RNA-seq profiling adult tissues from 42 echinoderm specimens from 24 orders and 37 families (1). The RNA-seq data were assembled, and translated into protein sequences. The *de novo* transcriptome assembly consisted of 1,198,706 amino acid sequences across 42 species. These data were clustered using OrthoMCL, an algorithm for grouping orthologous protein sequences based on sequence similarity (11). The resulting orthoclusters database consisted of groups of 749,397 orthologous and paralogous transcripts. The orthocluster data have been annotated through sequence similarity with respect to a purple sea urchin *Strongylocentrotus purpuratus* genome which was the best annotated echinoderm genome at the origin of the project (12). These annotations provide the basis for keyword searches.

The EchinoDB web-based application was initially built in the GO programming language in 2016 (1). The updated EchinoDB has been rewritten in R Shiny (3), which is highly extensible, easy to write, and maintain compared to the previous implementation. R Shiny supports faster development of user interfaces by providing a framework that requires no knowledge of scripting languages like HTML, CSS or JavaScript. In addition, the new version facilitates the inclusion of new data from collaborations. We have taken advantage of this feature to extend the application’s capabilities to expose new data such as the *Lytechinus* (6) and *Ophioderma* (5) sequences to interface for BLAST searches via Sequenceserver (4).

## Materials and Methods

In the latest EchinoDB release, we added a text box that allows users to conduct searches using accession numbers and other keywords with or without the use of wildcard entries. Results include protein sequence(s), annotated description(s), known NCBI GenInfo Identifier (GI ids), and orthocluster(s). The annotations are borrowed from aligning our sequences to the well-characterized protein sequence dataset of *Strongylocentrotus purpuratus* (i.e., sequences attributed to taxon 7668 in NCBI’s RefSeq, accessed in August 2012). These results can be further filtered by name or GenInfo Identifier (GI ids) in the search box in the top right corner. Additionally, users are able to expand or narrow their search based on taxonomic class, order, and family via toggle switches. Figure 1 depicts the design created in R Shiny for the EchinoDB application. Each row of the result table represents an orthocluster with the sequence similarity count or total hits. The number of hits is clickable, facilitating the viewing and downloading of related amino acid and nucleotide sequences in a FASTA format.

**Figure 1:**
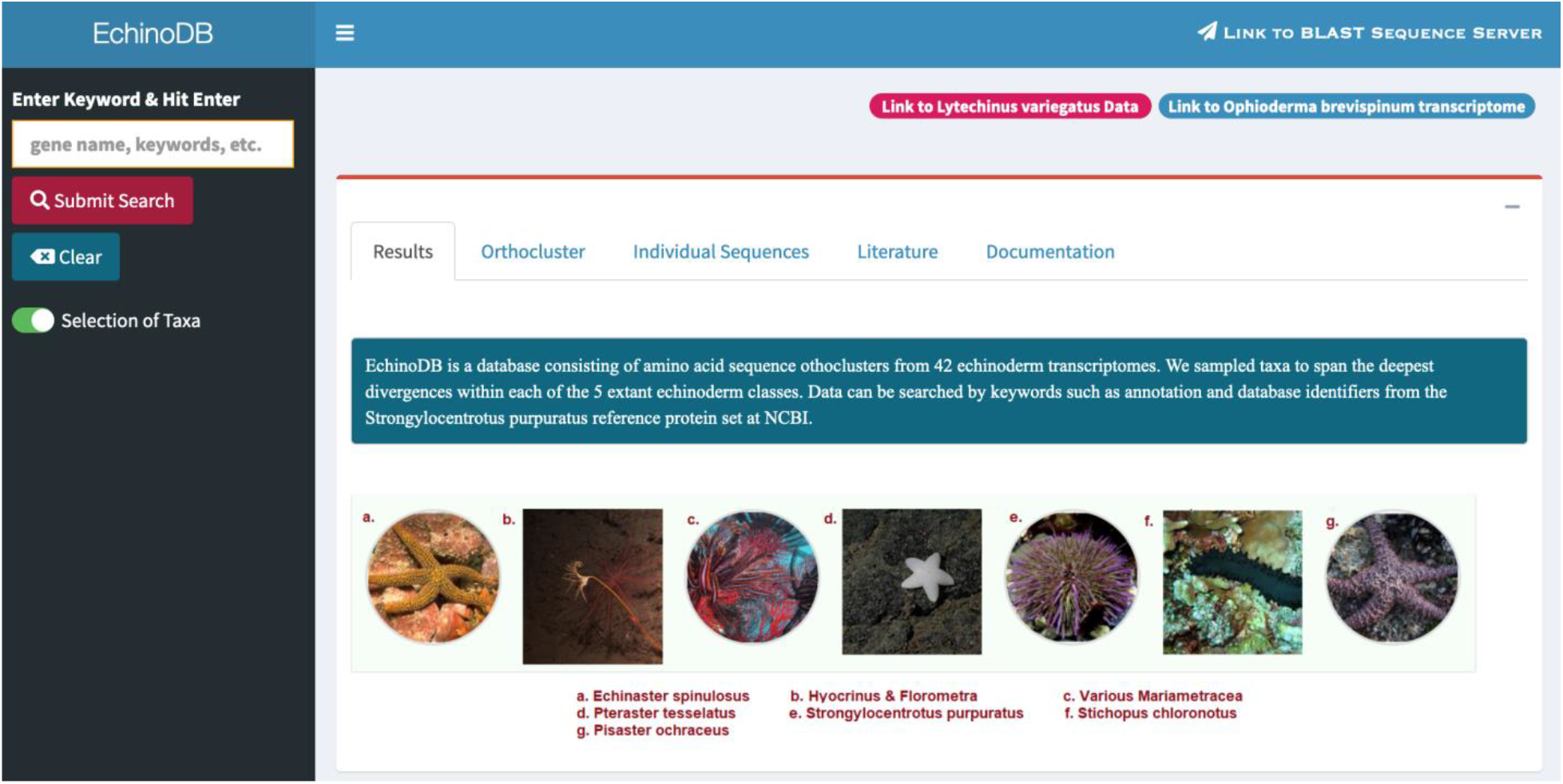
Screenshot of our EchinoDB landing page (19), available at https://echinodb.uncc.edu. Users could search against all extant echinoderm classes, orders, and families or un-toggle to retrieve information for a particular taxon.

### New data resources for ophiuroid and echinoid

We have added RNA-seq data of *O. brevispinum* (5), a common brittle star found in shallow waters of the western Atlantic Ocean ranging from Canada to Venezuela. We have also added high-quality genome assembly data of *L. variegatus* (6), a sea urchin found in shallow waters throughout the western Atlantic Ocean ranging from the United States to Venezuela.

#### OphiuroidDB

We have provided a transcriptome dataset, translated and annotated using BLASTX functionality against the NCBI collection of predicted proteins of *S. purpuratus* (25) and protein models from UniProt’s Swiss-Prot (16) and NCBI’s RefSeq (17). The application can be accessed via “Link to *O. brevispinum* transcriptome” in EchinoDB. This page is referred to as “OphiuroidDB” and it serves the data exclusive to the brittle star, *O. brevispinum* (20).

These data were first used to characterize the downstream genes controlled by the Notch signaling pathway, which is believed to play a role in brittle star regeneration (5). Then, the raw sequencing reads were submitted to the NCBI as a GEO dataset under the accession number GSE142391 (5, HYPERLINK \l "bookmark26" 24), and transcriptome sequences can be downloaded from OphiuroidDB. A total of 30,149 genes were identified and included in the application.

#### EchinoidDB

EchinoidDB facilitates access to a recently published annotated high-quality chromosomal-scale genome assembly of *L. variegatus* (6, HYPERLINK \l "bookmark23" 21). The data *(Lvar_3*.*0)* includes 27,232 nucleotide and protein sequences which were annotated using BLASTP (18) against UniProt Swiss-Prot (16), *S. purpuratus* (23) and non-*S. purpuratus* RefSeq invertebrate protein models (17). These annotations can be downloaded from EchinoidDB.

### Use case example

To demonstrate the usage of EchinoDB and associated resources, we can search for genes associated with the Notch signaling pathway. This should serve as a relevant example because this pathway is required for regeneration in echinoderms (5). Knowledge of the Notch signaling pathway is important because it is highly conserved in the animal kingdom and regulates various cellular events, including proliferation, differentiation, fate specification, and cell death (26-29).

For example, the user can search for Notch-related genes and obtain the corresponding sequences and metadata from our web resources. To do this, the user can search for the keyword Notch in our web resources to locate Notch-related sequences in brittle stars and other echinoderms. The results include NCBI’s accession numbers, other unique identifiers, descriptions, start and end positions, and other details depending on the application used. In addition, the interface allows the user to visualize any selected sequence or cluster of sequences from the related repository.

## Results and Discussion

EchinoDB is designed to facilitate comparative transcriptomic studies of deeply-sampled clades of echinoderms (1). EchinoDB is re-factored in R Shiny. R Shiny is extensible and code developed with R Shiny can be integrated, readily with CSS themes, HTML widgets, and scripting languages like JavaScript. In addition, as R Shiny is widely adopted, the code can be modified and tuned at later stages in the development cycle by many developers.

EchinoDB is hosted using the Nginx web server (2) in Amazon Web Services (AWS) (7). AWS offers on-demand cloud computing services to build your own web-based applications independent of university information technology bureaus.

### Application features

As many echinoderm “omic” data are not yet well annotated, blast search is an important complement to keyword or accession search.

#### Using Sequenceserver to run BLAST

EchinoDB contains an instance of Sequenceserver (4), a web-based BLAST server that supports sequence similarity searches against nucleotide and protein sequence databases. EchinoDB provides nucleotide and protein databases to facilitate sequence similarity searches using default or user-selected parameters.

Integration with BLAST allows users of EchinoDB to search data resources with strings of the query sequence. Figure 2 illustrates Sequenceserver for BLAST functionality and can be accessed via “Link to BLAST Sequenceserver” in the EchinoDB application.

**Figure 2:**
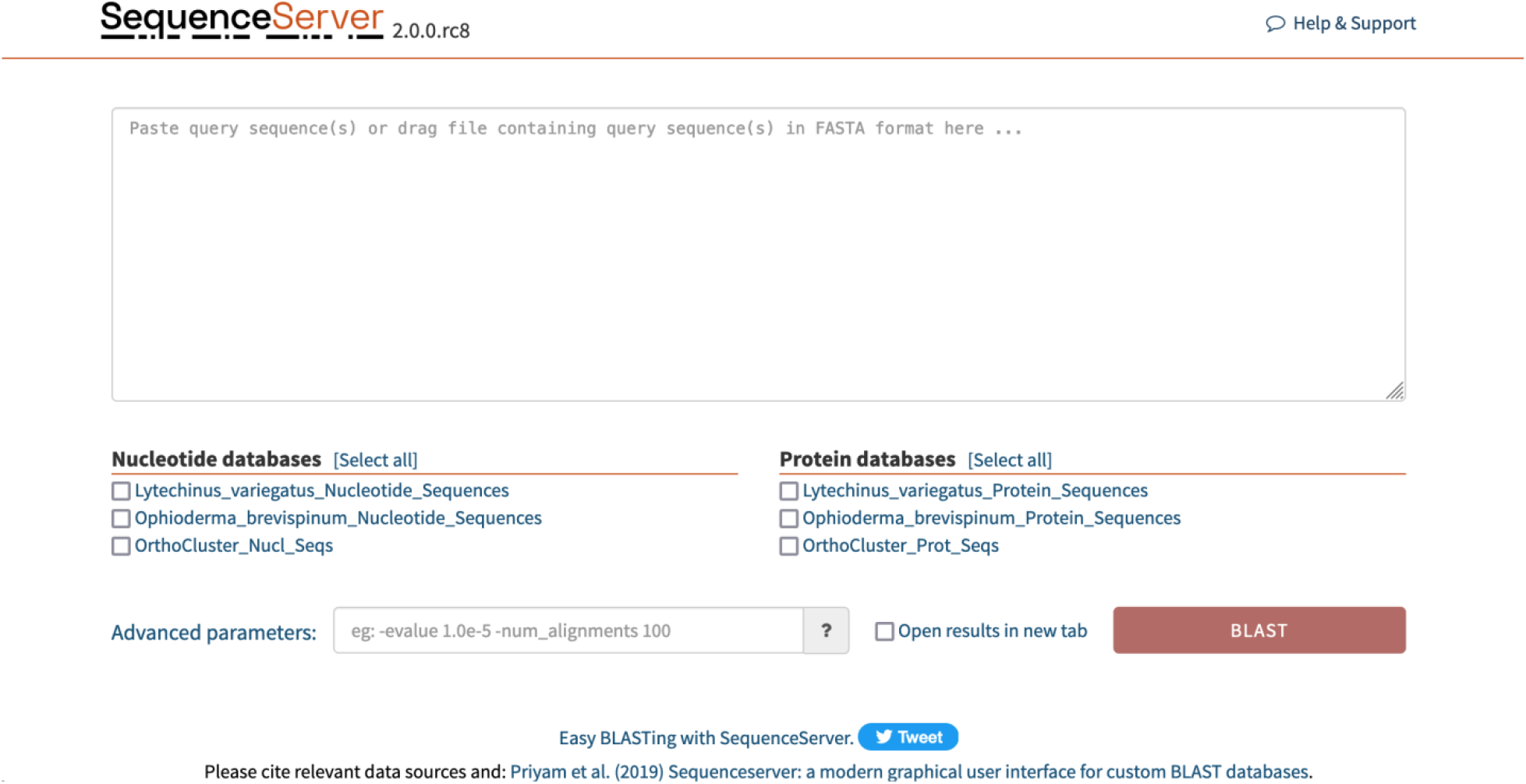
Screenshot of our Sequenceserver integrated in EchinoDB (22). Users can perform BLAST searches against nucleotide and protein sequences of included datasets in the application via https://echinodb.uncc.edu/sequenceserver/.

#### Literature

We provide a repository that contains links to the research papers associated with EchinoDB by their title. The literature repository is updated regularly.

#### Additional data

A link is added in the Literature section to allow users to download data. For example, one dataset provides evidence that *Xyloplax* sp. is a velatid (order of class Asteroidea) asteroid rather than a new class (8). The data included in EchinoDB includes tables and phylogenomic data from large amounts of transcriptome data used in this paper. The additional data repository is updated regularly.

### Usage and documentation

EchinoDB, EchinoidDB, and OphiuroidDB user manuals (Additional files 2–4: Files S1–3, respectively) are available in a tab named “Documentation” in the EchinoDB website. The user manuals are downloadable and provide instructions with screenshots to assist the user in navigating through the application.

Figure 3 demonstrates the usefulness of EchinoDB and some of its outputs. The figure depicts the usage in a step-by-step process by locating individual or clusters of Notch-related amino acid sequences in brittle stars and other echinoderms.

**Figure 3:**
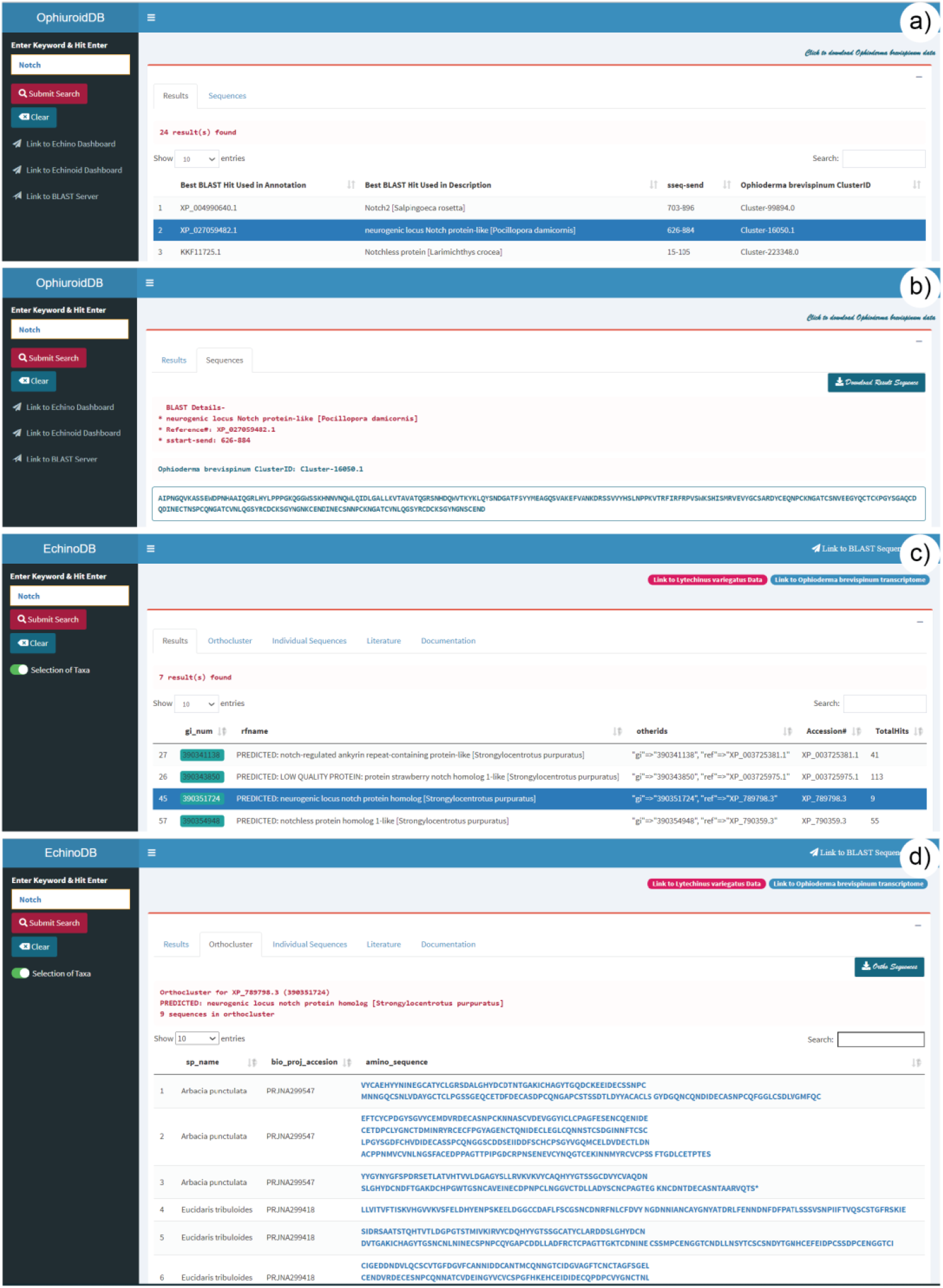
Usage example illustrating the search for Notch-related sequences in brittle stars and other echinoderms. **a)** Screenshot of the OphiuroidDB main page (https://echinodb.uncc.edu/BStarApp/). The image shows the results after searching for the keyword “Notch” against the database of the brittle star *Ophioderma brevispinum*. The interface allows the selection of any record on the results page to view the sequence. **b)** Example amino acid sequence from one selected Notch-related gene in OphiuroidDB. **c)** Results after searching for the keyword “Notch” in EchinoDB (https://echinodb.uncc.edu). In this example, the search was conducted against the repository of clusters of orthologous genes discovered from echinoderm transcriptomes. A selected record will be highlighted, and amino acid sequences from the Orthocluster repository will be displayed. d) Displays amino acid sequence clusters of the selected orthologous record of the Notch-related gene from the EchinoDB repository.

## Conclusion

The updated EchinoDB provides, via a cloud-based server, additional tools and data from collaborations and our lab that can be of interest to a variety of scientific communities. One of our focal points in the future is to extend the genomic, transcriptomic, and orthocluster contents of EchinoDB.

## Supplementary Materials

Assembled sequences and orthoclusters are available in EchinoDB (https://echinodb.uncc.edu; 19). Raw reads from the various echinoderm species are available in NCBI’s SRA (see accession numbers in Additional file 1: Table S1). Additionally, the user manuals for EchinoDB, EchinoidDB, and OphiuroidDB are available as Additional file 2: File S1, Additional file 3: File S2, and Additional file 4: File S3, respectively. Additional files are available in Zenodo (https://doi.org/10.5281/zenodo.5803547).

## Acknowledgements

Research reported in this publication was supported by the National Institute of General Medical Sciences of the National Institutes of Health under award number R15 GM128066-01. We acknowledge the support of several entities of the University of North Carolina at Charlotte including the College of Computing and Informatics, the Graduate School, the Department of Bioinformatics and Genomics, University Research Computing, and the Bioinformatics Research Center. Additionally, we thank Benjamin Stalcup for issuing a SSL certificate for EchinoDB application and Steven Blanchard for helping us with certificates deployments and instructions for web server setup. We also thank Shantoy Hansel and Jan Kofsky for their testing of the application. Finally, we thank Greg Wray for providing the genomic data of the green sea urchin, *Lytechinus variegatus* (6).

## Notes

https://doi.org/10.5281/zenodo.5803547

### Competing Interest Statement

The authors have declared no competing interest.

