## Supplemental File S1 for "EchinoDB: An update to the web-based application for genomic and transcriptomic data on echinoderms"

### EchinoDB User Manual

**Author:** Varnika Mittal,
**Date:** Nov. 1, 2021

#### 1. Access EchinoDB Application

Click on the following link to access the application: https://echinodb.uncc.edu

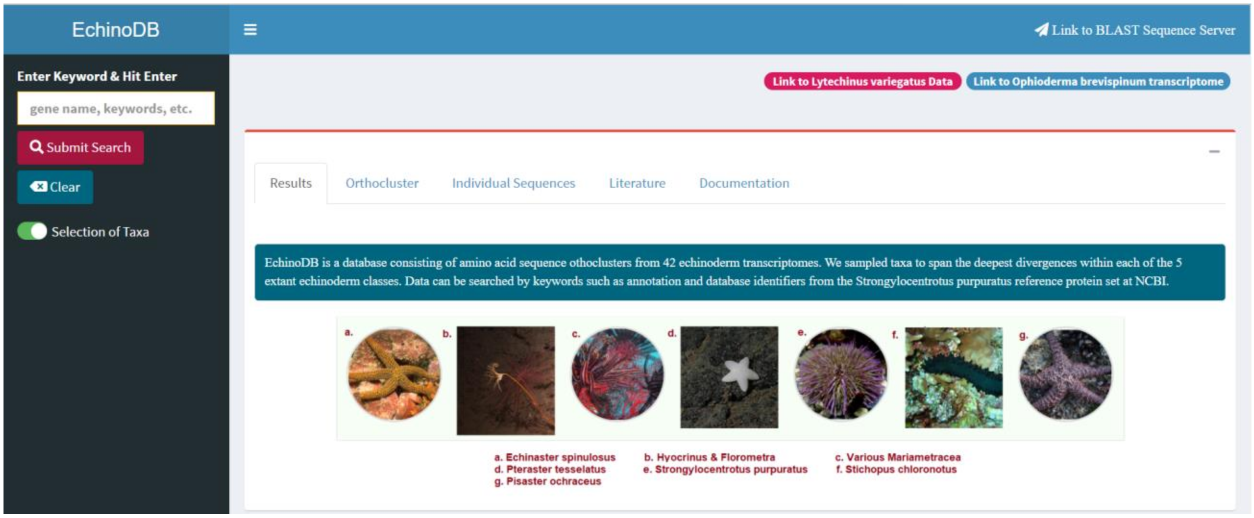


#### 2. Perform Search against Orthocluster Database

Type in any any keyword such as “zinc,” “chlor,” “iron” or NCBI’s accession numbers like “XP_78042.” Hit the “Submit Search” button in the upper left side of the web page or press the “Enter” key to execute search.

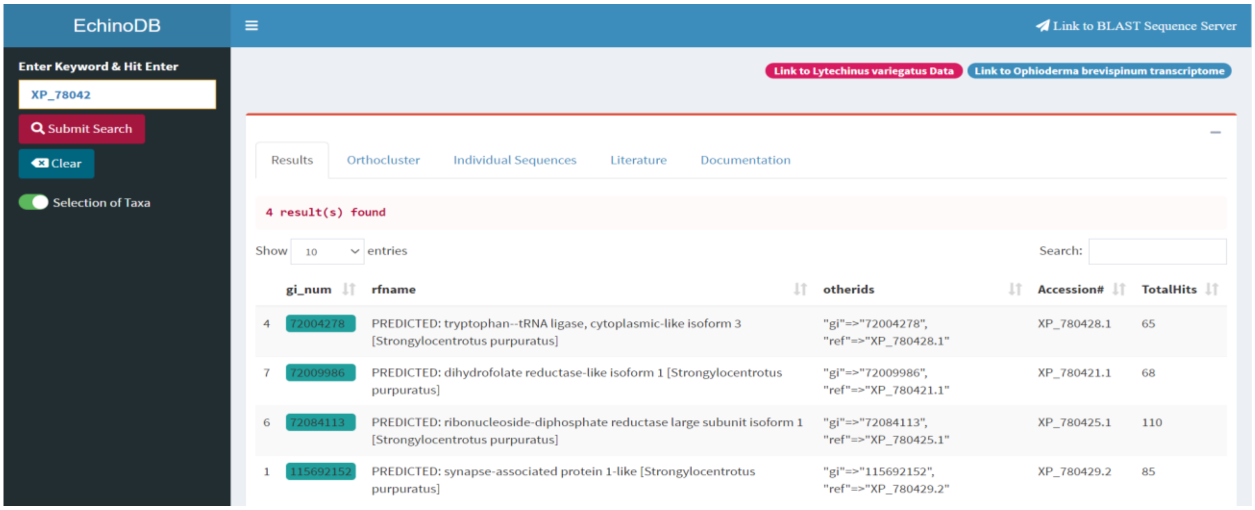

➢ Note that, using the accession number XP_78042, 4 results are found in this example. However, using the zinc keyword, 98 results are found. These numbers may change as we update the database.

- **Filter Results**
  All the names extant Echinoderm classes, orders, and families for which we have data can be used as keywords to filter the search. If information to be retrieved for particular taxon, un-toggle "Selection of Taxa" in the left pane.

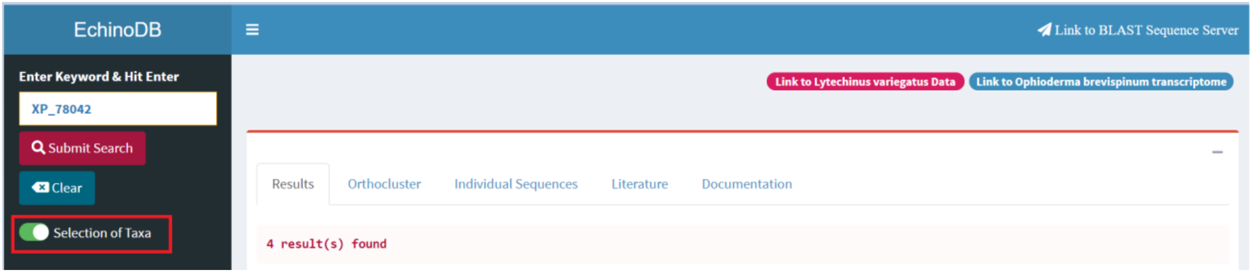


➢ Select Asteroidea and hit the submit button to conduct a search within this class of echinoderms.

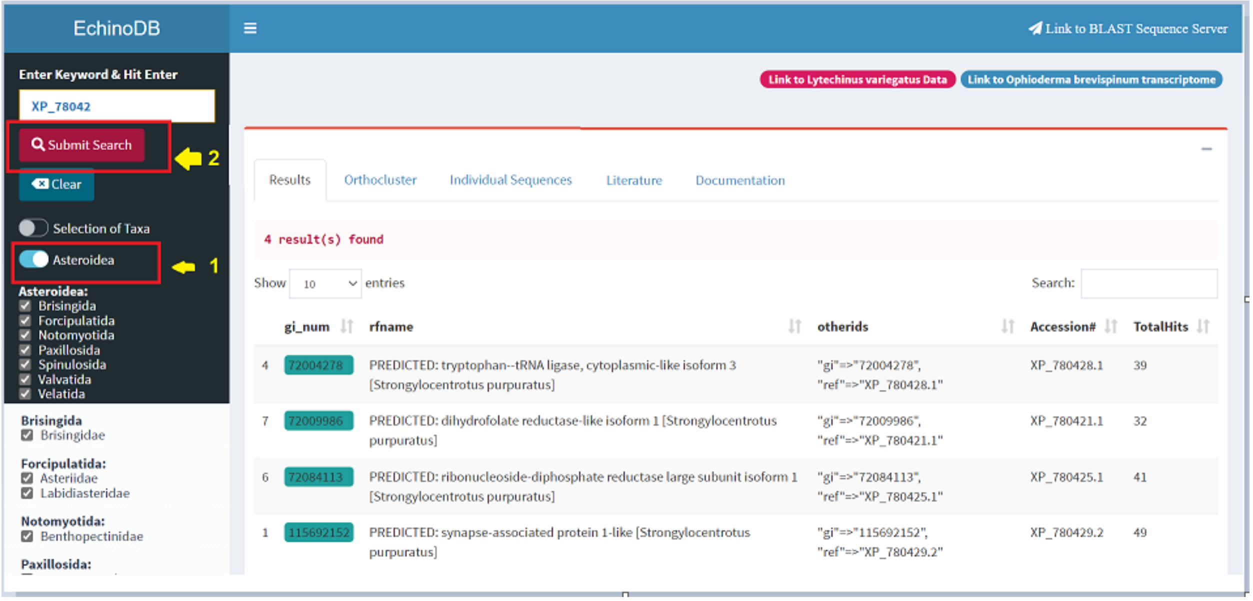

➢ Unselect families to include only those which you would like to use in your search.

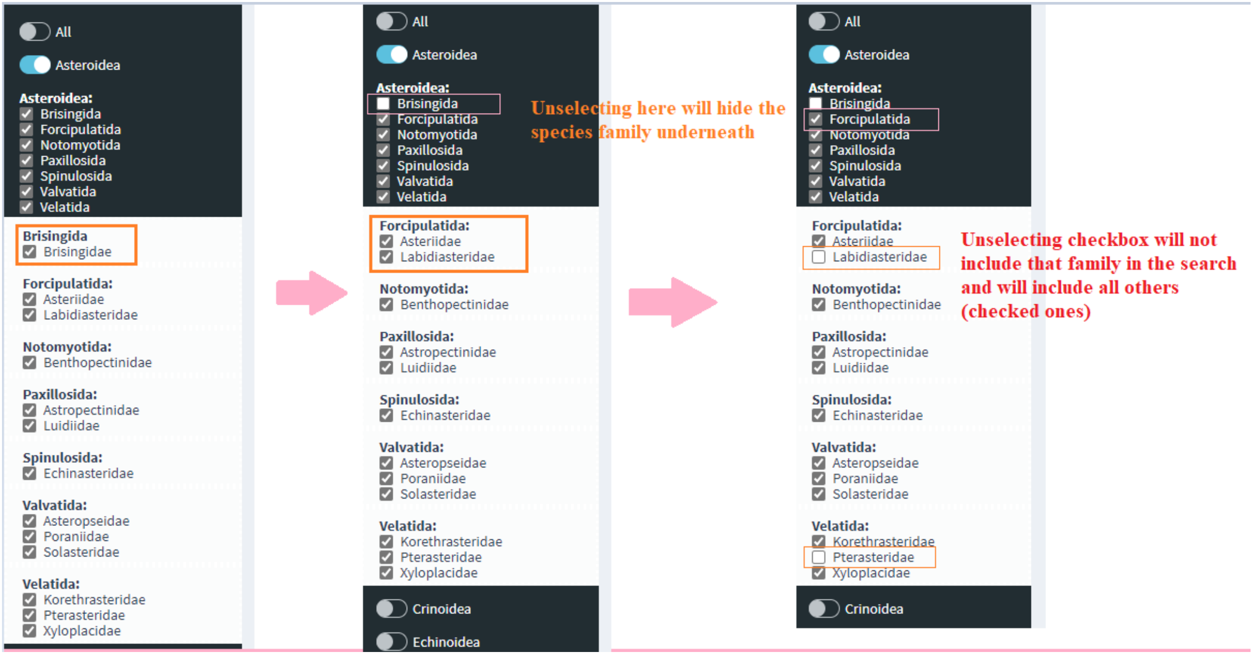


➢ Hit the search button with all these filters applied to narrow down the results.

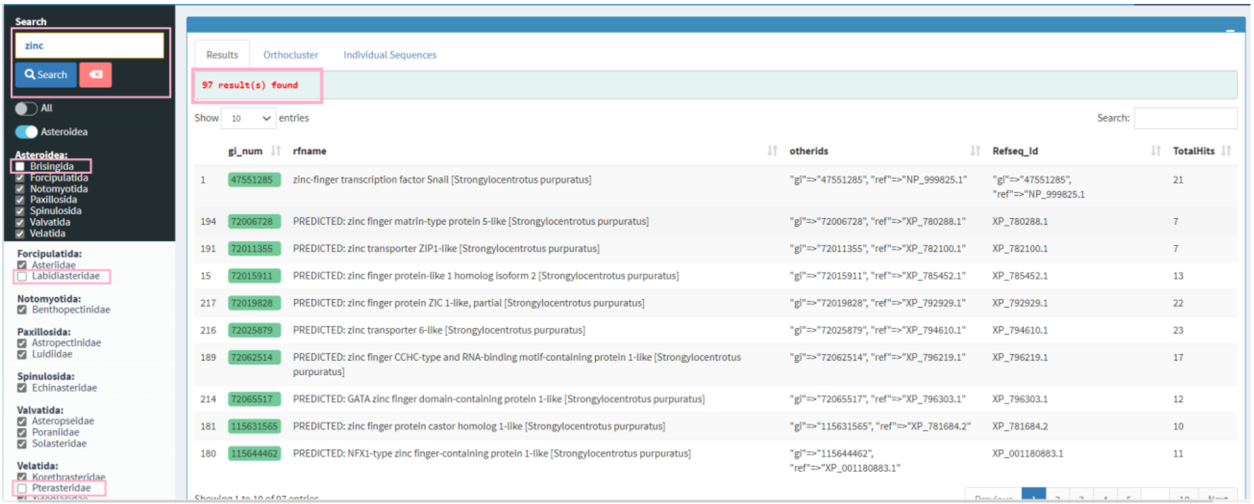

- **Search Based on Taxonomic Group**
  The user can narrow the search to, for example, only a single family of echinoderms. In the example below we are checking only the family Solasteridae within the order Valvatida of the class Asteroidea.

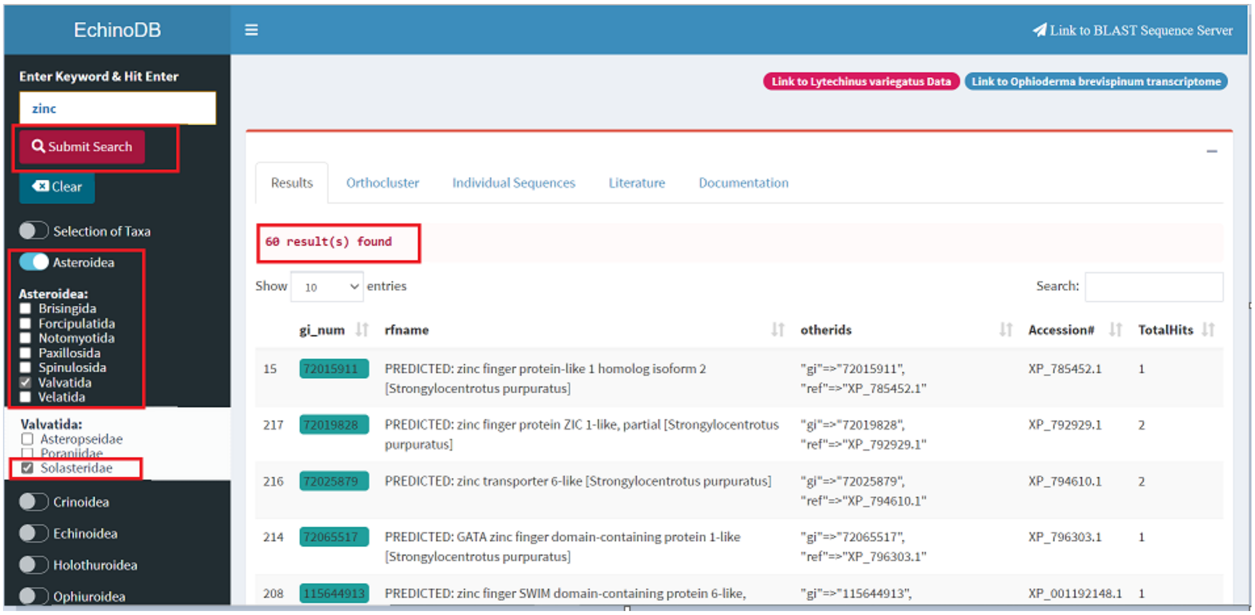


- **Other Search Options**
  You can search by name or gi number in the search box in top right corner.

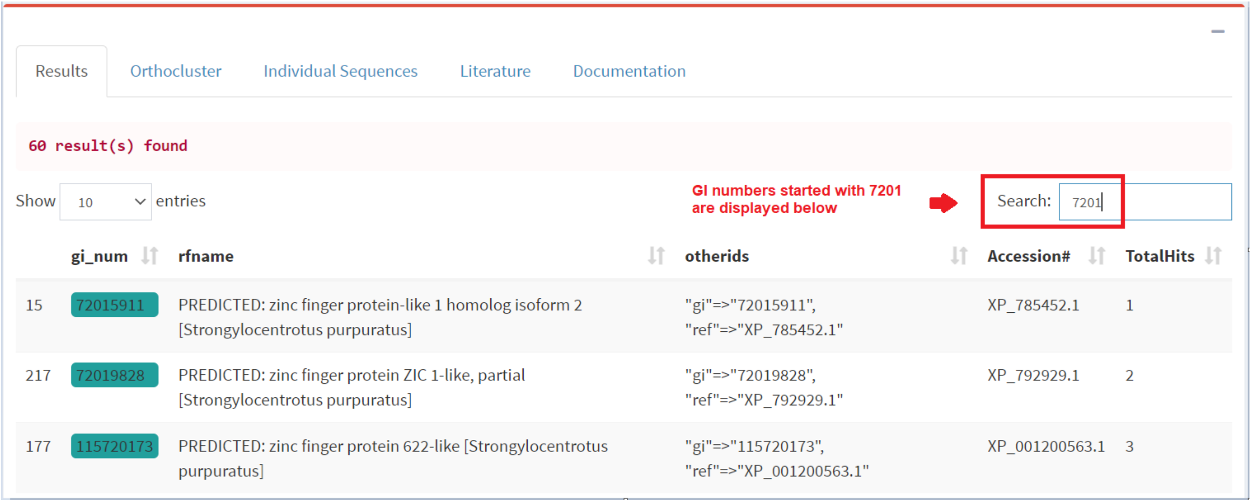

#### 3. View Orthocluster Sequences

Select a whole row and the record will be highlighted in blue (you can select only one row at a time).

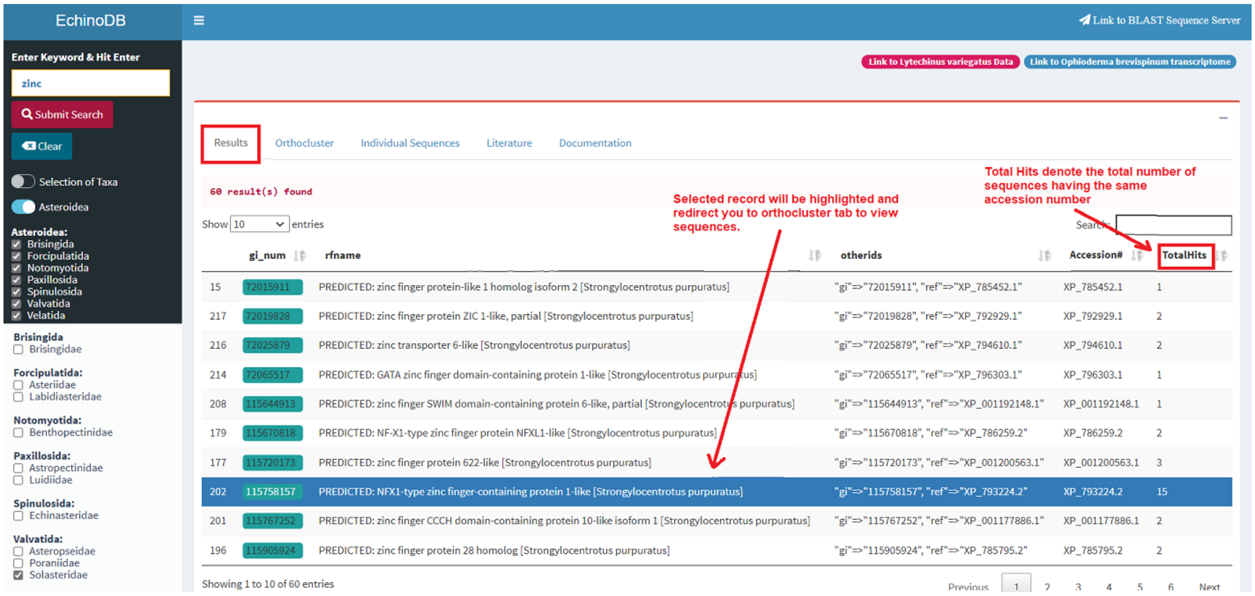

- **Amino Acid Sequences**
  Display amino acid sequences from the selected orthocluster.

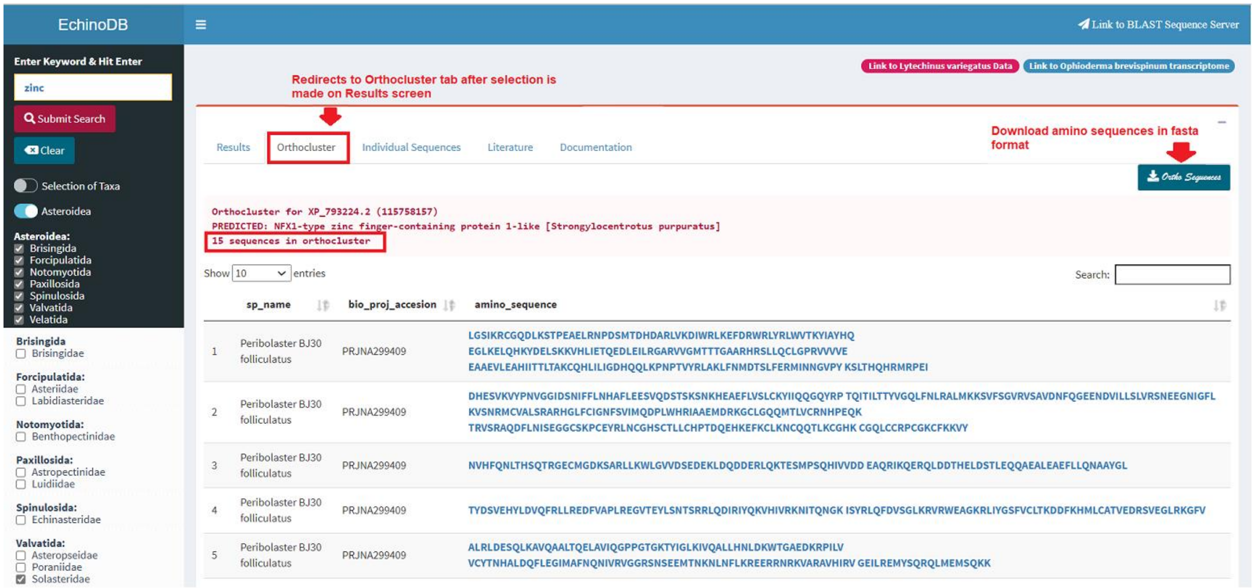

- **Individual Sequences Tab**
  Select a row in Orthocluster results which contain amino acids sequences to further view related nucleotide sequences.

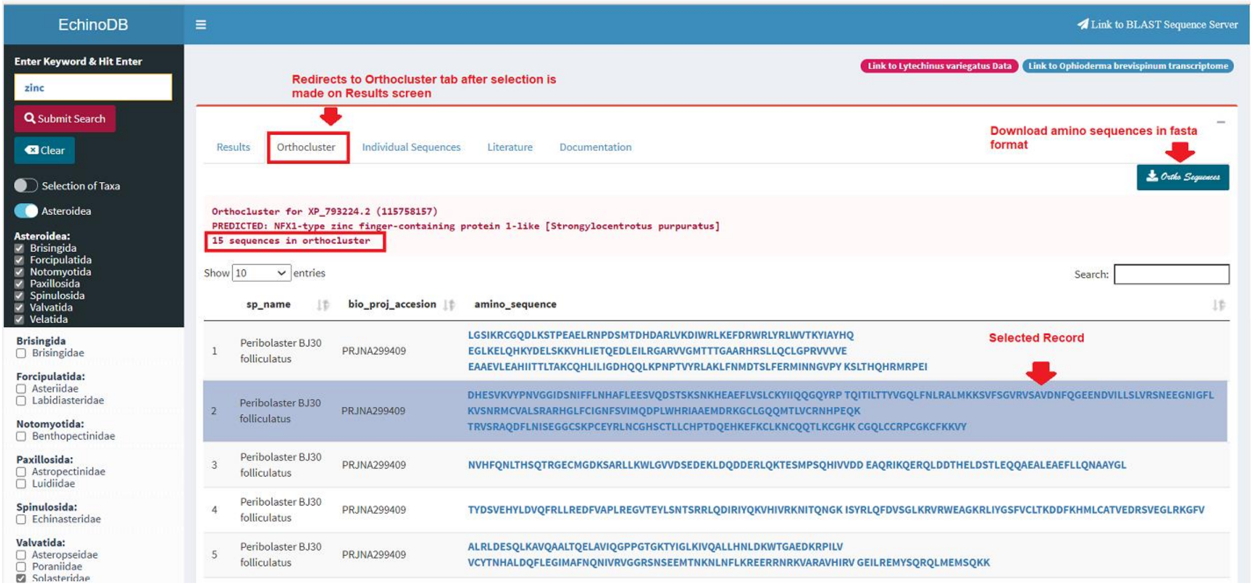


- **Amino Acid and DNA Sequences**
  The user will find different buttons to download amino acid (protein) or DNA sequences.

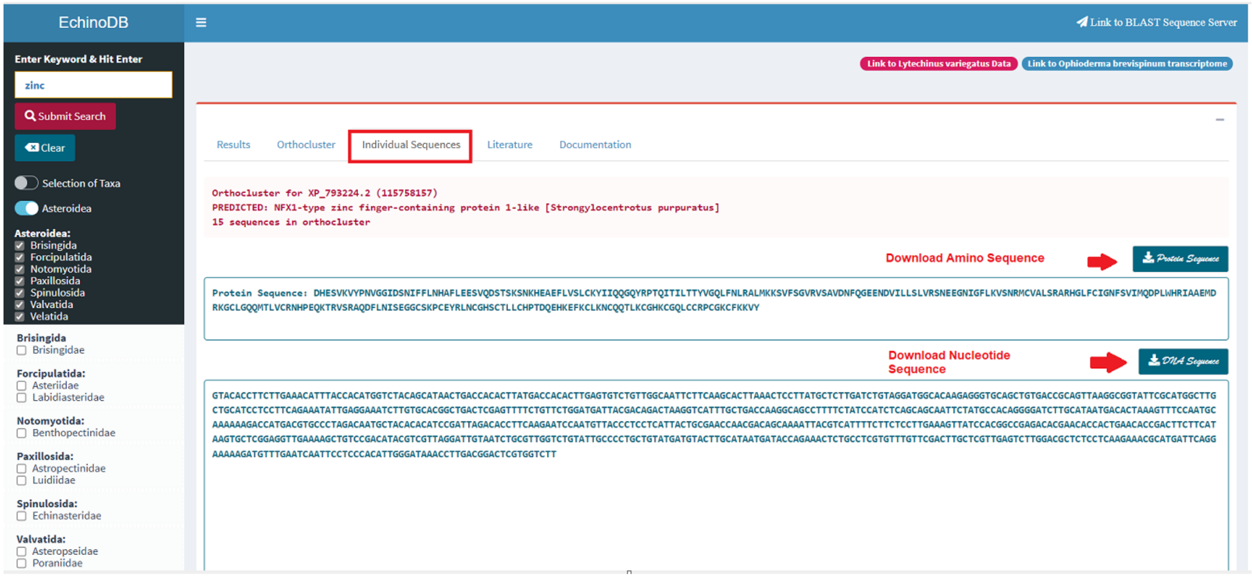

#### 4. Sequenceserver for Basic Local Alignment Search Tool

Access Sequenceserver (Priyam *et al*. 2019) from EchinoDB by clicking the “Link to Sequenceserver” in the header line or by type in the following URL: https://echinodb.uncc.edu/sequenceserver

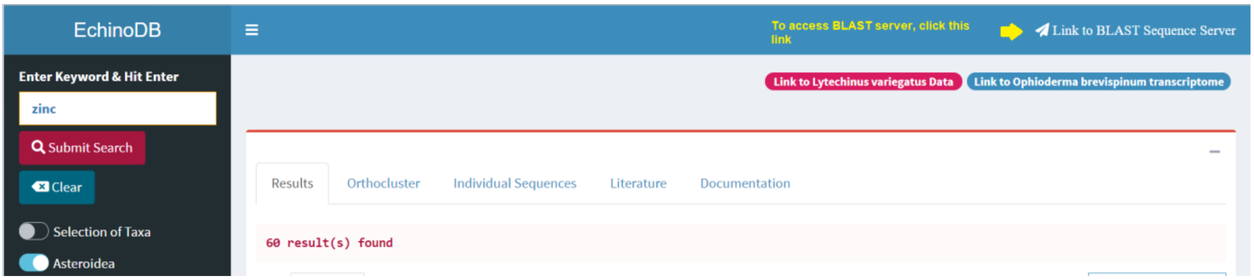


- ➢ Paste your query string (amino acid or nucleotide sequences) in the text area to perform BLAST search.

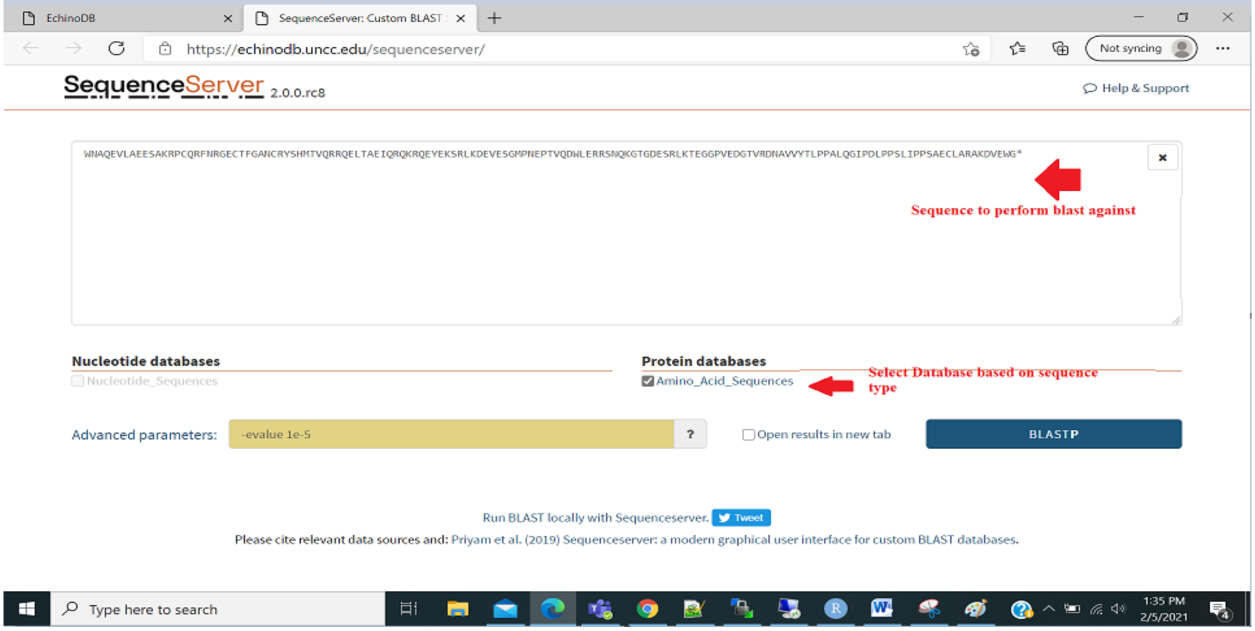

- **Sequenceserver Results**

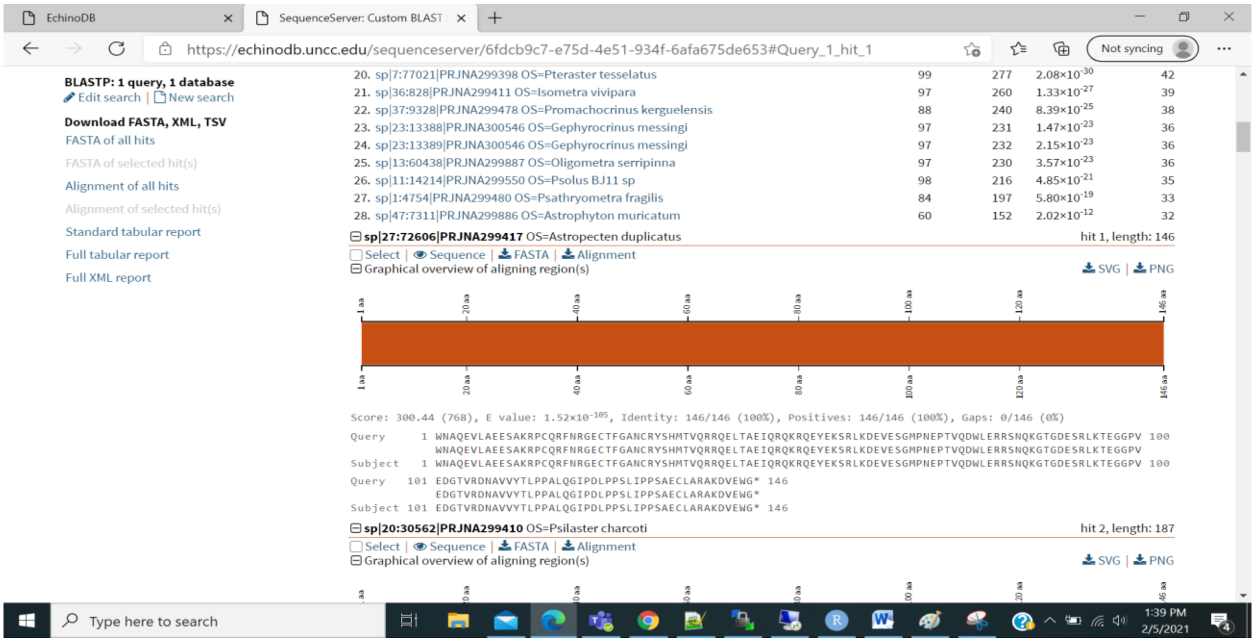

- **Download FASTA Sequence from a high scoring pair from Sequenceserver**

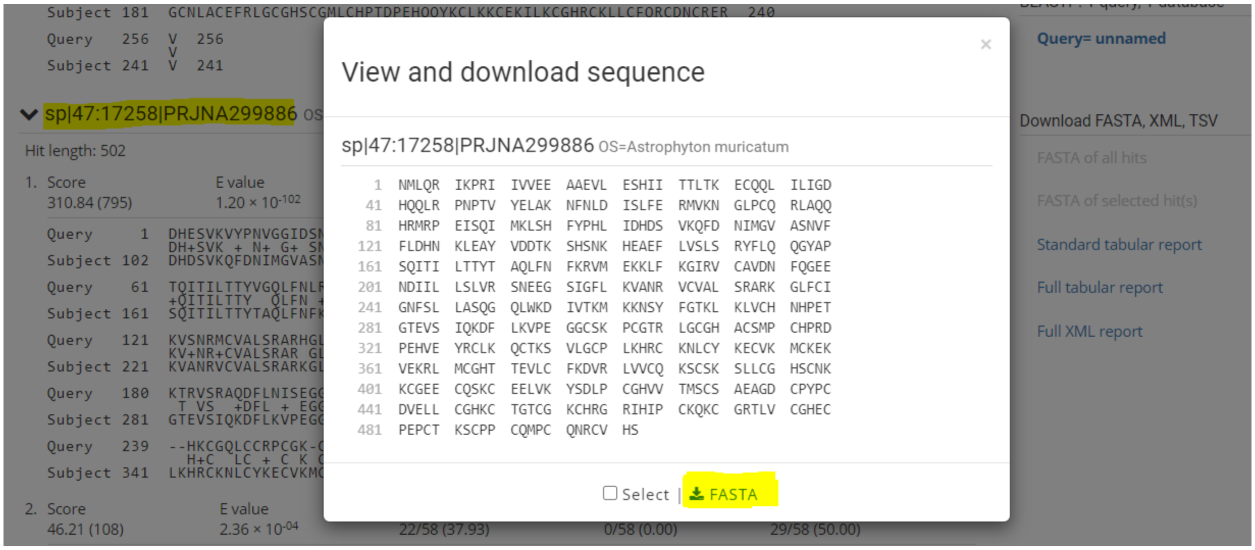

#### 5. Clear Search/Results

Go to the results tab and hit the “Clear” button underneath the search button to clear search.

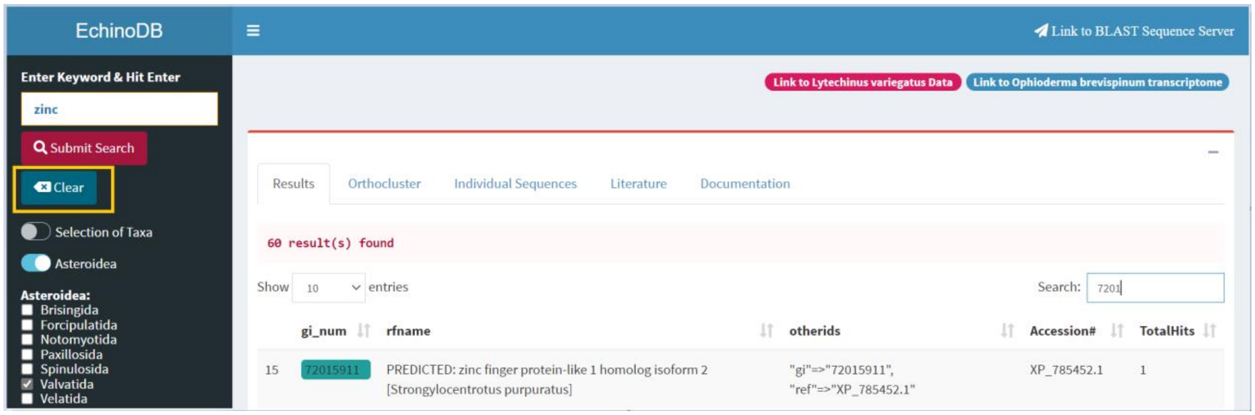


- The search will be cleared after the button is clicked. The user can turn the switch “Selection of Taxa” on if further taxon selection is desired.

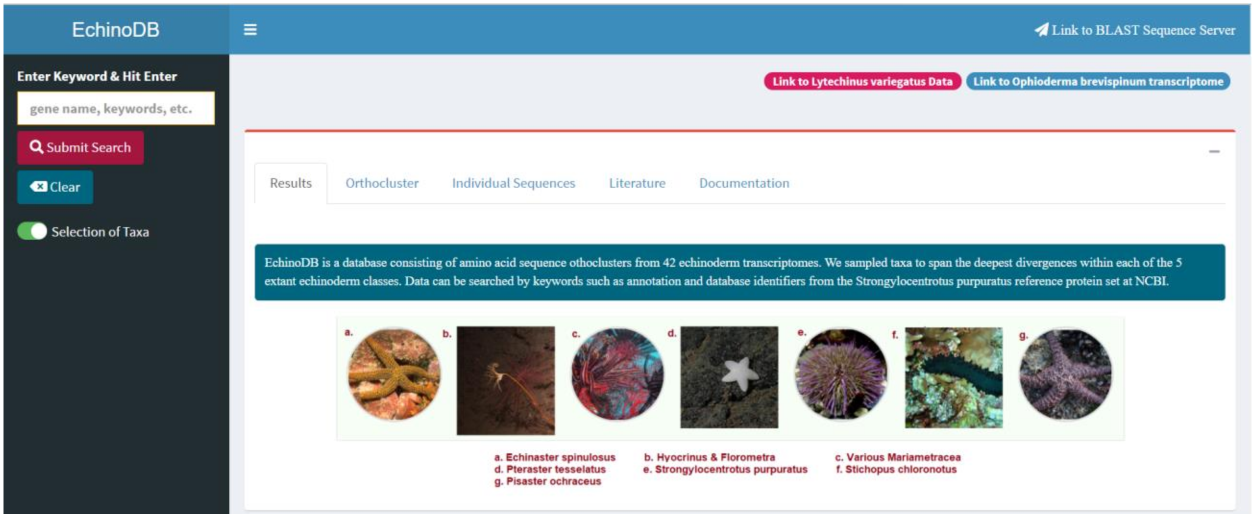

#### 6. Additional Links

Links in the top right are provided to redirect users to “OphiuroidDB” by clicking blue button (“Link to *Ophioderma brevispinum* transcriptome”) or “EchinoidDB” by clicking red button (“Link to *Lytechinus variegatus* Data”).

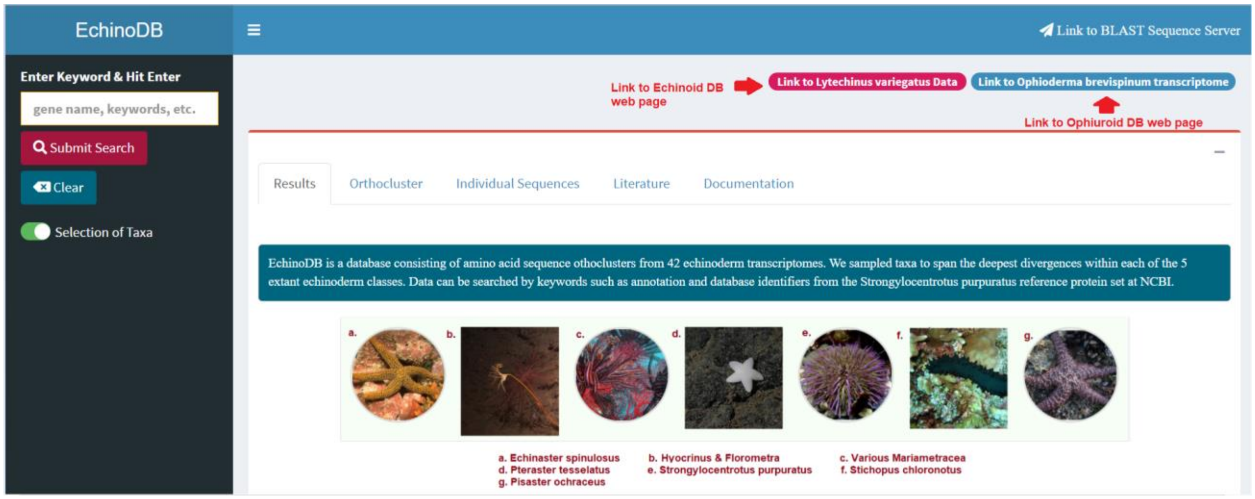
