## Supplemental File S2 for "EchinoDB: An update to the web-based application for genomic and transcriptomic data on echinoderms"

### Echinoid Database (EchinoidDB) User Manual

**Author:** Varnika Mittal,
**Date:** Nov. 2, 2021

#### 1. Access the EchinoidDB Application

Click on the following link to access the application: https://echinodb.uncc.edu/SUrchinApp

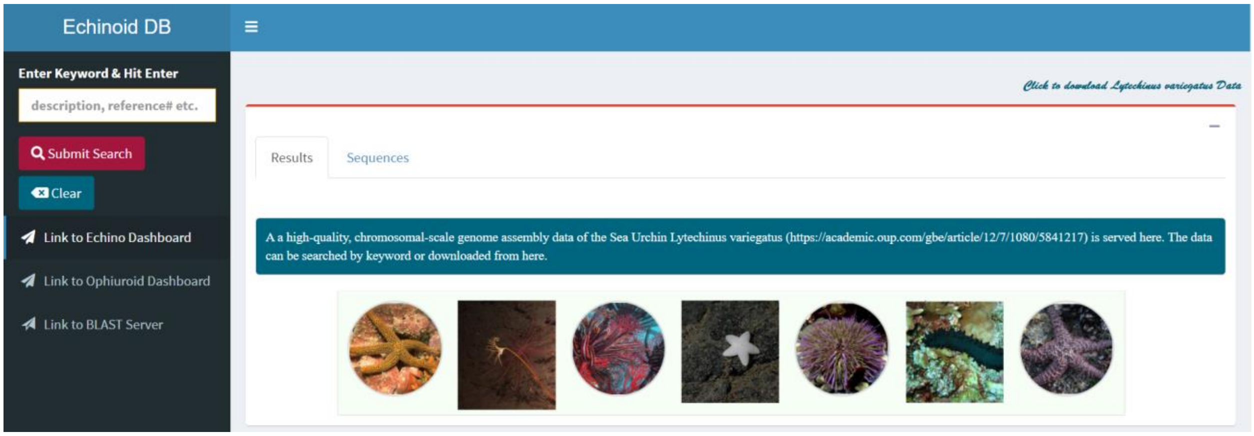

#### 2. Search Within the Genome of the Sea Urchin *Lytechinus variegatus*

The user can search data in the chromosome-level assembly of the sea urchin *Lytechinus variegatus* (Davidson et al. 2020). Use any keyword such as “zinc”, “chlor”, “iron” or an NCBI’s accession number such as XP_0220792. Hit the “Submit Search” button in the upper left side of the web page or press “Enter.”

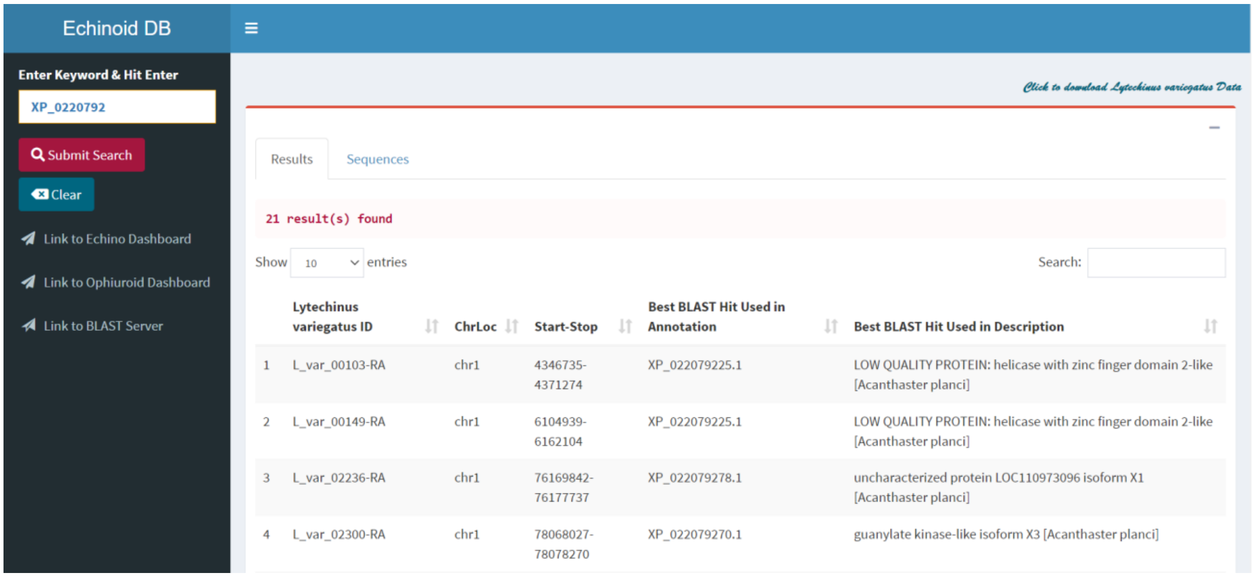


- ➢ In the example above we used the accession number XP_0220792 keyword and found 21 results. Note that the results may change as we update our databases.

- **Other Search Options**
  You can search by description or NCBI’s accession number in the search box in top right corner.

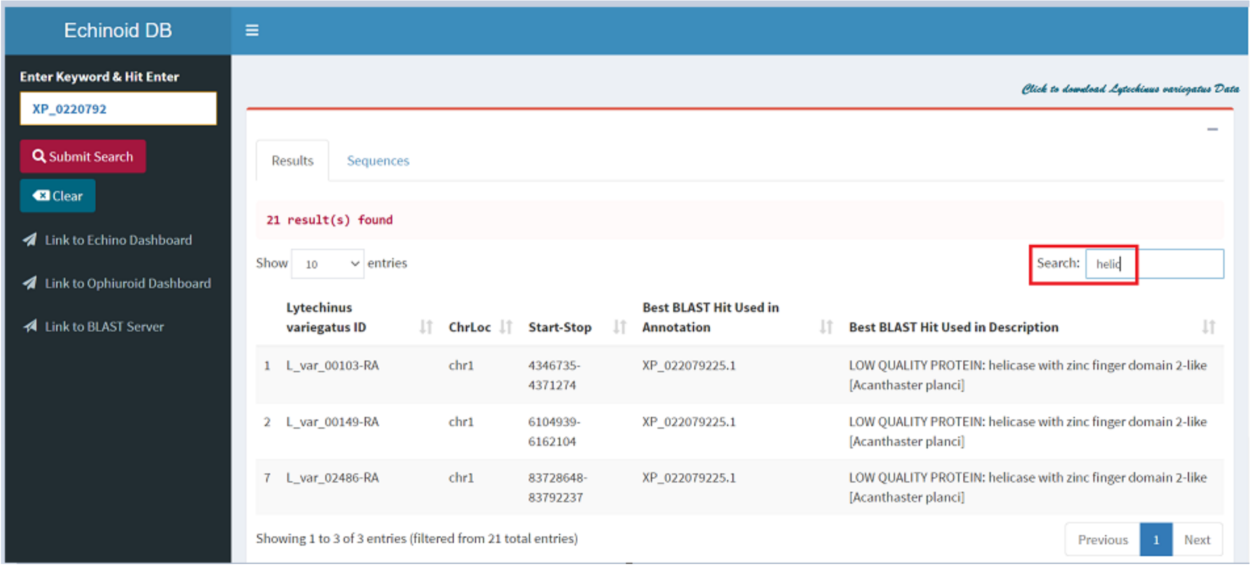

- **Download Data**
  The link “Click to Download *Lytechinus variegatus* Data” is provided in the top right corner to download *Lytechinus variegatus* transcriptome sequences.

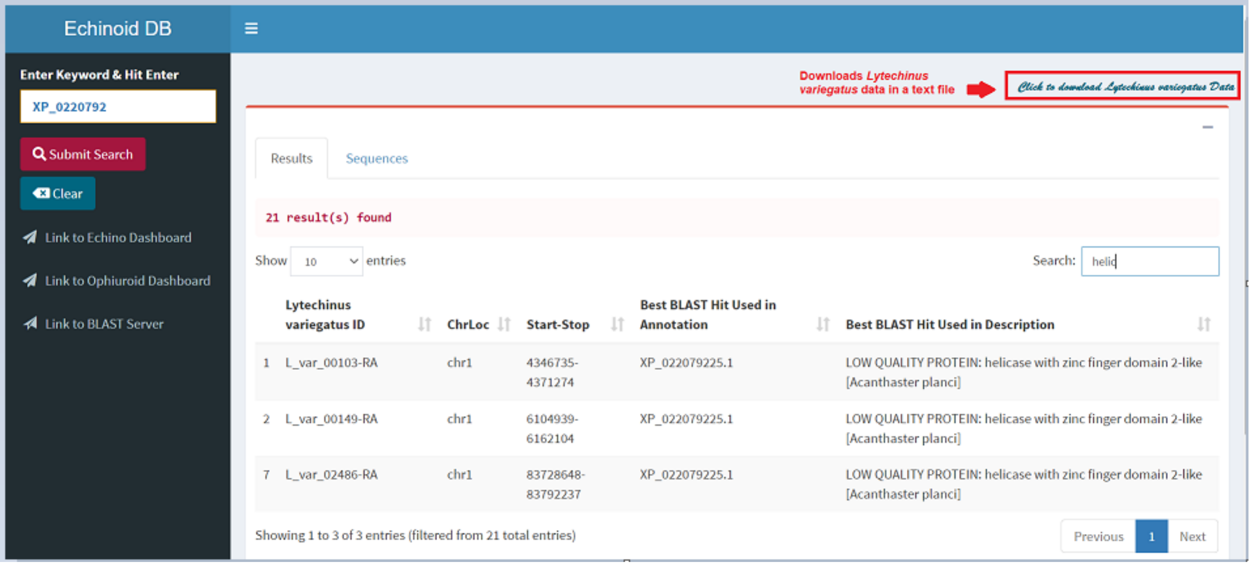

#### 3. Visualize *Lytechinus variegatus* Sequences

Select a whole row and the record will be highlighted in blue (the user can select only one row at a time).

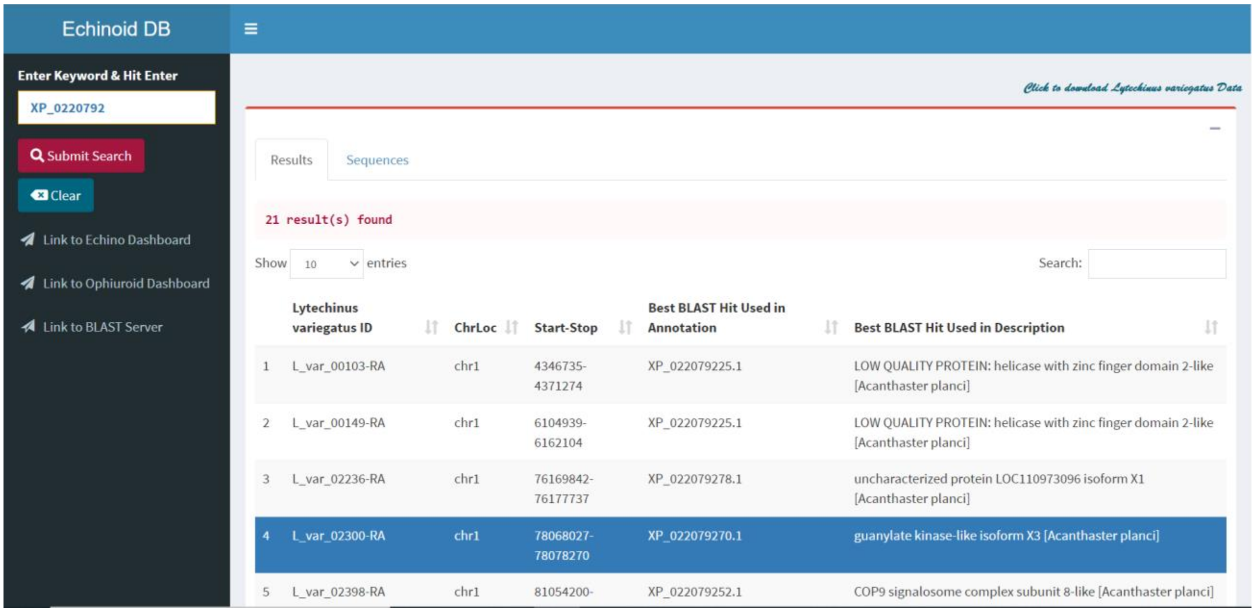


- **Sequences Tab**

  After the record is selected, the user is redirects to the “Sequences” tab and display protein sequences from Ophiuroids repository. Furthermore, it allows downloading the search results in FASTA format.

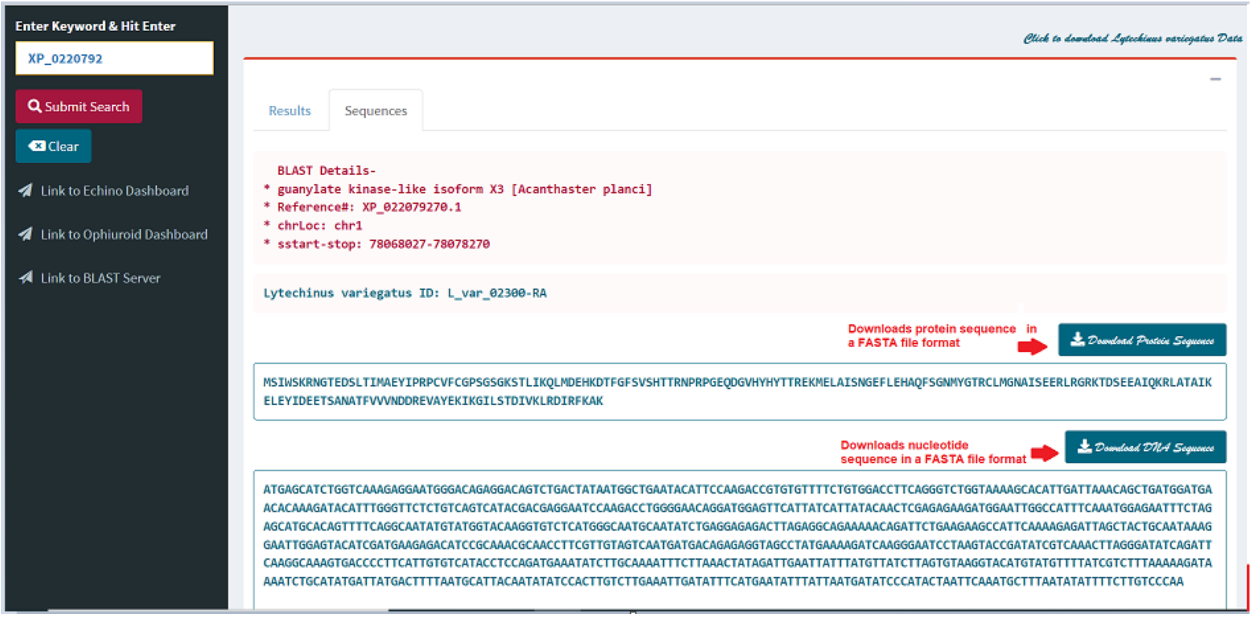

#### 4. Using Sequenceserver

The user can access Sequenceserver (Priyam et al. 2019) to run BLAST by clicking the "Link to BLAST Server" in the left pane or by typing in the following URL: https://echinodb.uncc.edu/sequenceserver/

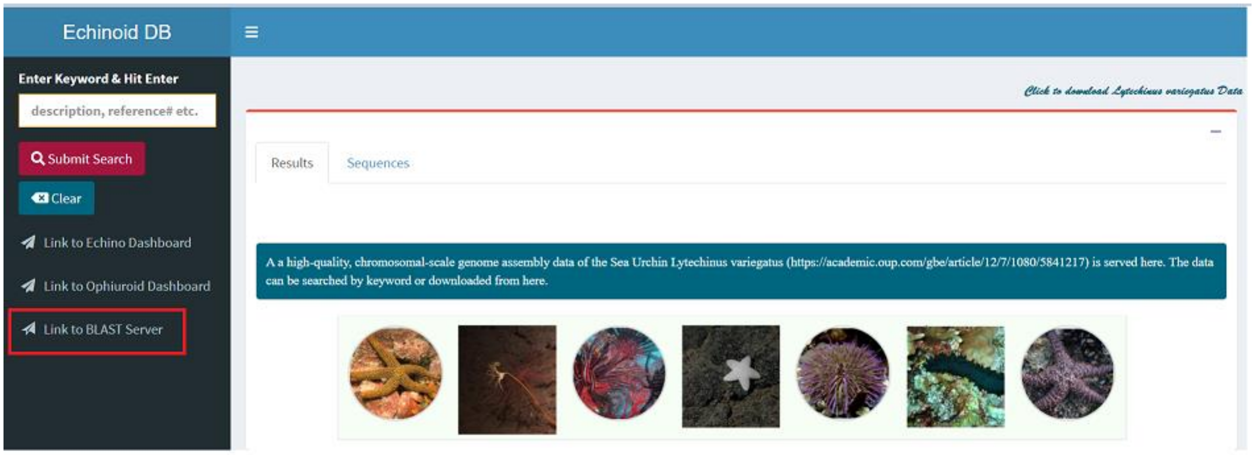


- ➢ Paste your query string (amino acid or nucleotide sequences) in the text area to perform a BLAST search.

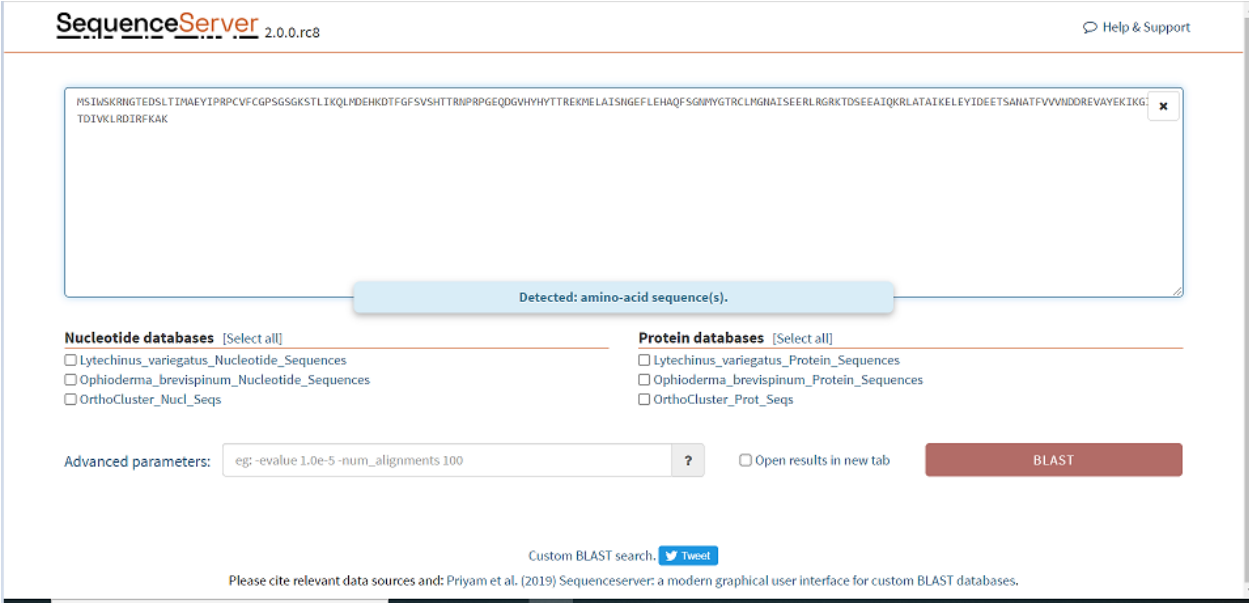

➢ Select database(s) to perform a BLAST search against the query sequences.

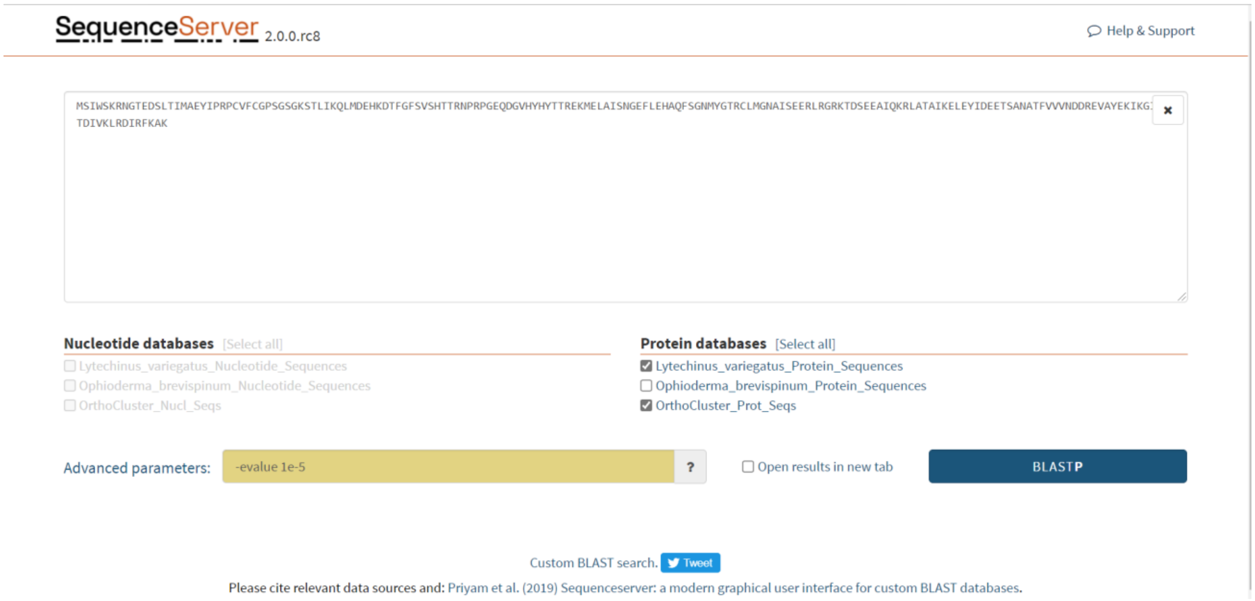


- **Sequenceserver Results**

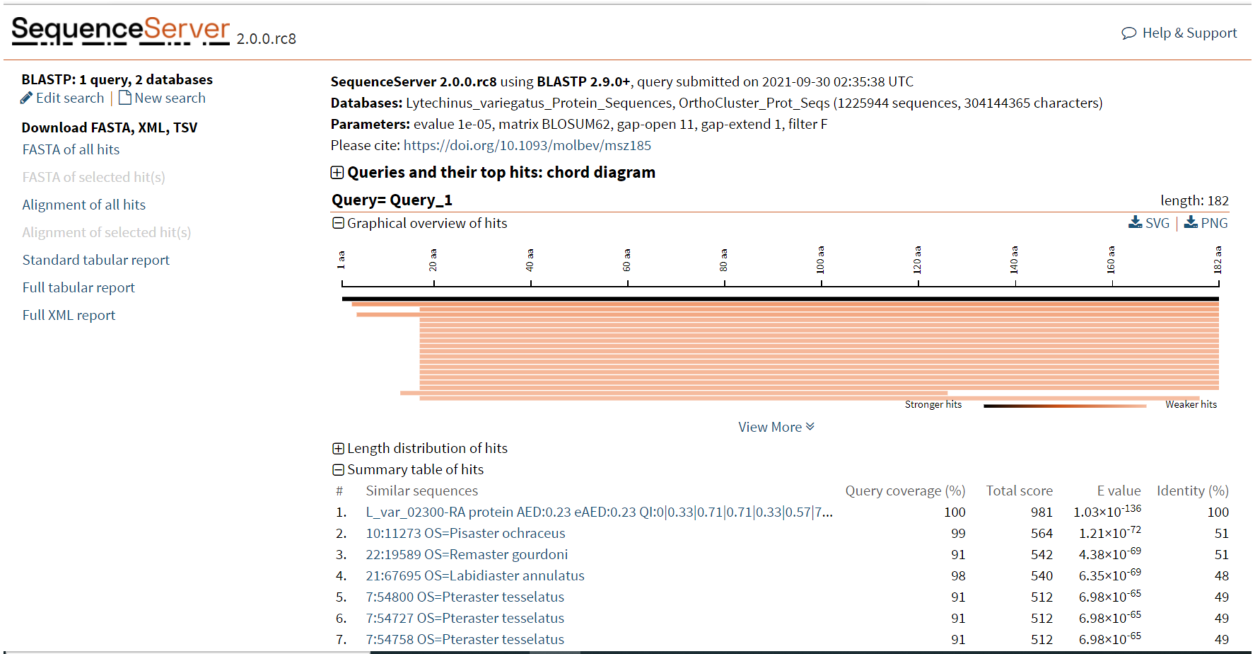

- **Download Sequenceserver Results**
  The user can use the buttons “FASTA” and “Alignment” to download data corresponsing to the results sequences or alignments, respectively.

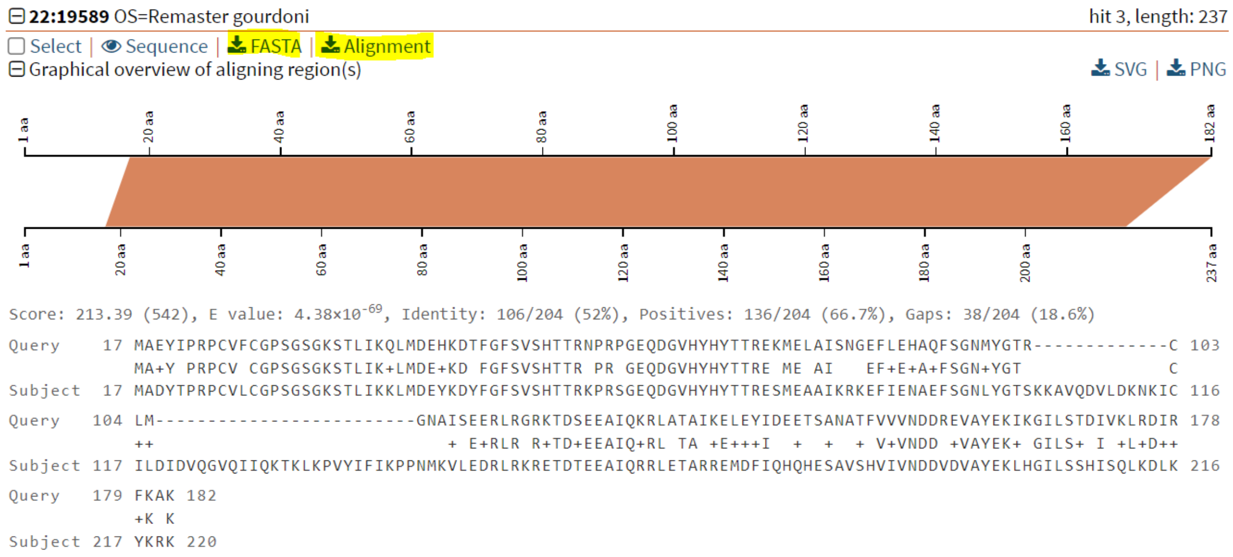

#### 5. Clear Search/Results in OphiuroidDB

Go to the results tab and hit “Clear” button to clear the search. Alternatively, the user can clear the search results by pressing “Delete.”

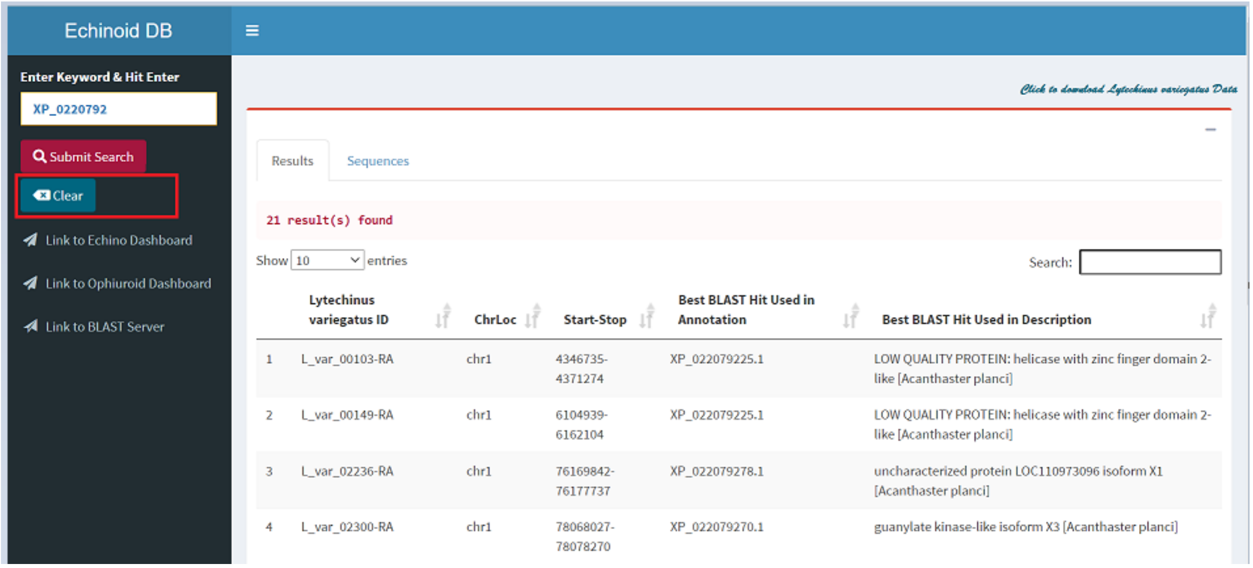

➢ The search will be cleared after the button is clicked or delete key is pressed.

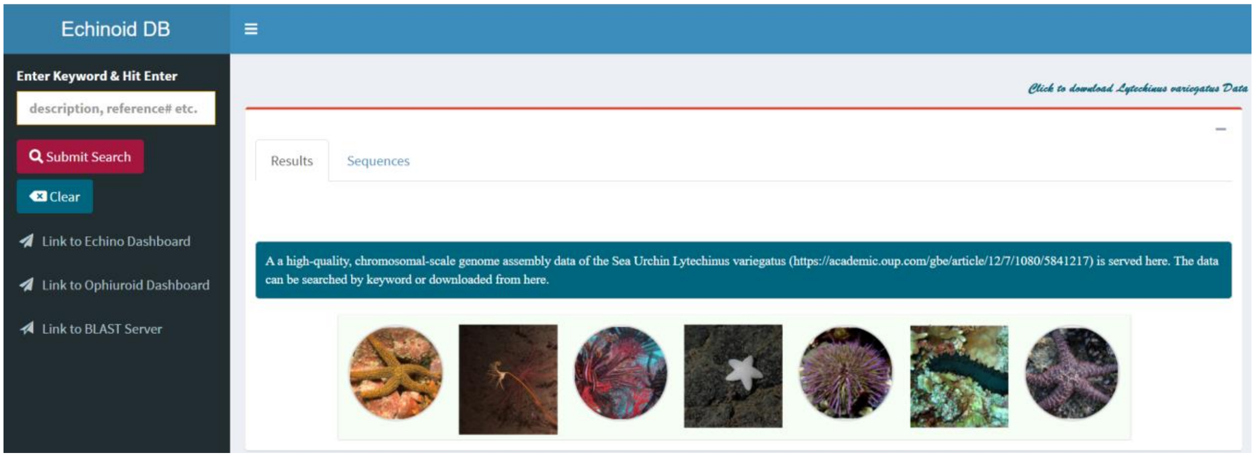


#### 6. Additional Links

Links in the left pane are provided to redirect users to "EchinoDB" or "OphiuroidDB" page.

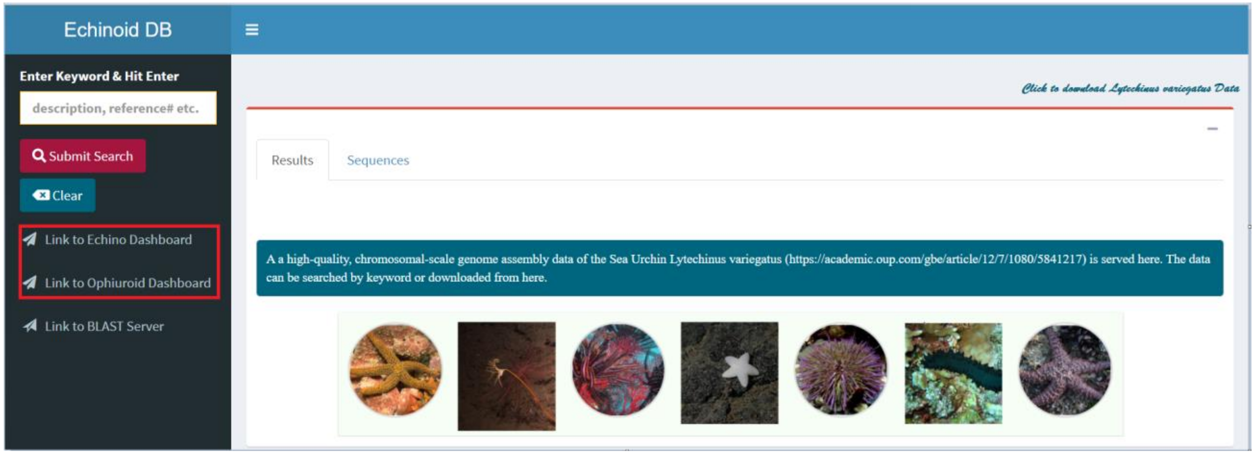
