## Supplemental File S3 for "EchinoDB: An update to the web-based application for genomic and transcriptomic data on echinoderms"

### Ophiuroid Database (OphiuroidDB) User Manual

**Author:** Varnika Mittal,
**Date:** Nov. 2, 2021

#### 1. Access Ophiuroid DB Application

Click on the following link to access the application: https://echinodb.uncc.edu/BStarApp

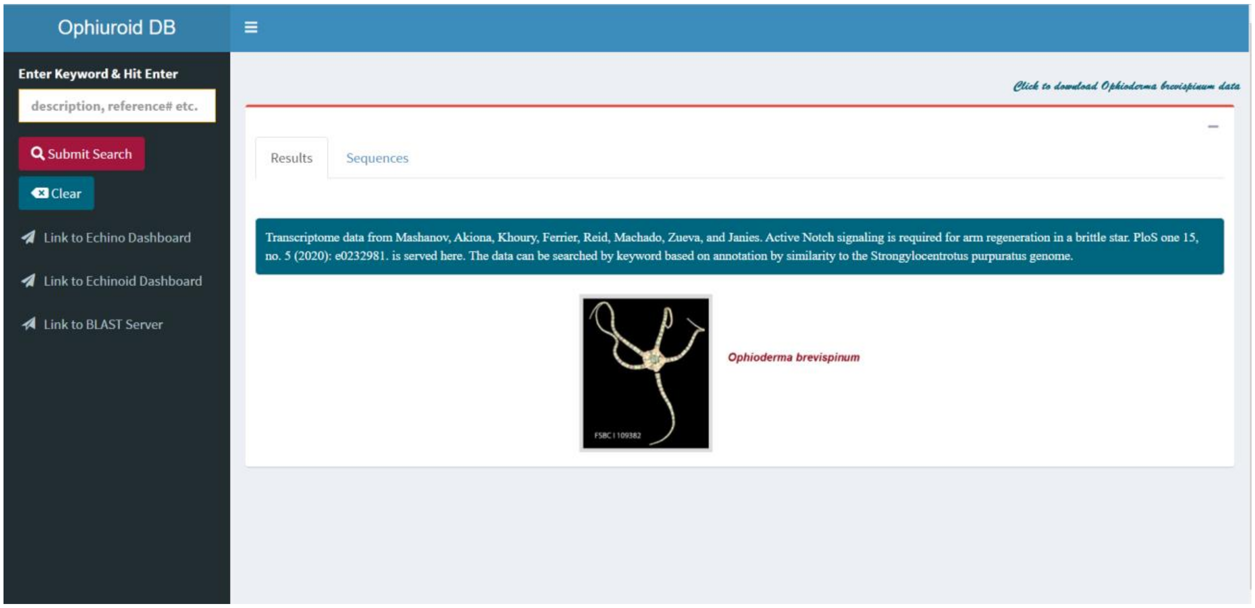

#### 2. Perform Search against Transcriptome Data for *Ophioderma brevispinum* in Ophiuroid Database (Mashanov et al. 2020)

The user can search the transcriptome data of the brittle star *Ophioderma brevispinum* (Mashanov et al. 2020) in the Ophiuroid Database. Use any keyword or string such as “zinc”, “chlor”, “iron” or NCBI’s Reference Number such as XP_0221120 for conducting search. Hit “Submit Search” button or “Enter” key in the upper left side of the web page.

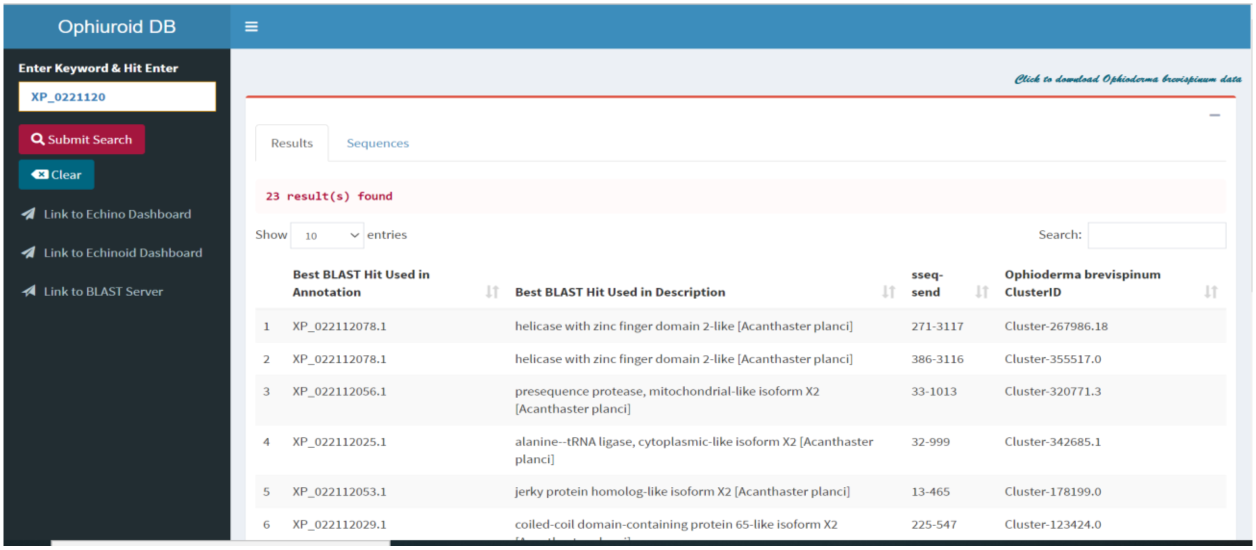


➢ With XP_0221120 keyword, 23 results are found.

- **Other Search Options**
  You can search by description or NCBI’s Reference Number in the search box in top right corner.

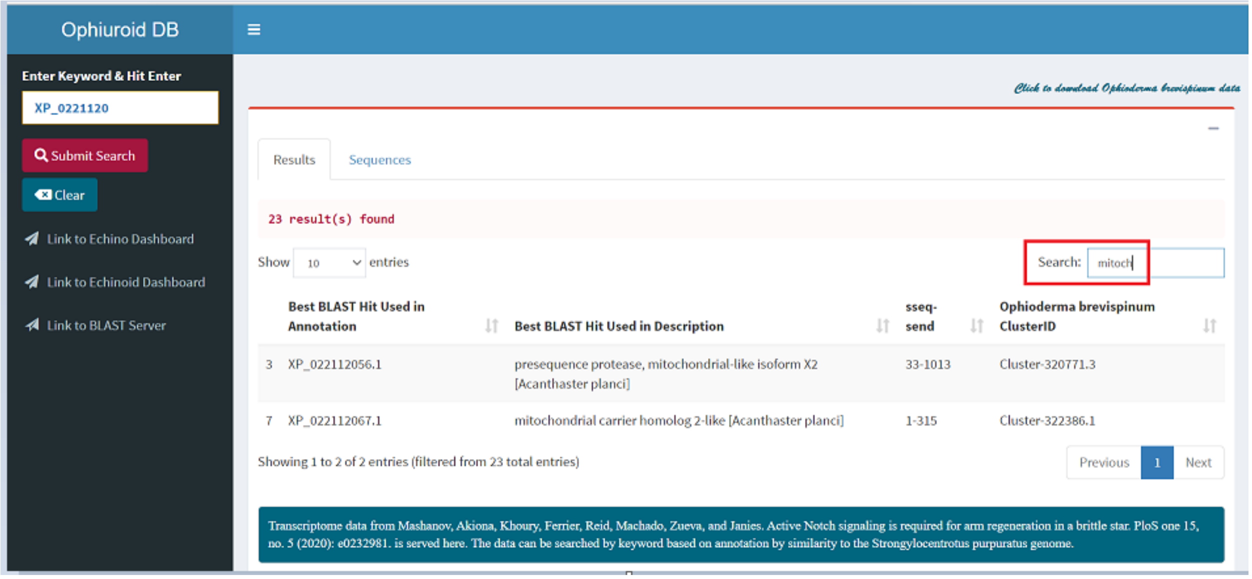

- **Downloading Search Results**
  A link “Click to download *Ophioderma brevispinum* data” is provided in the top right corner to download transcriptome sequences for *Ophioderma brevispinum* species.

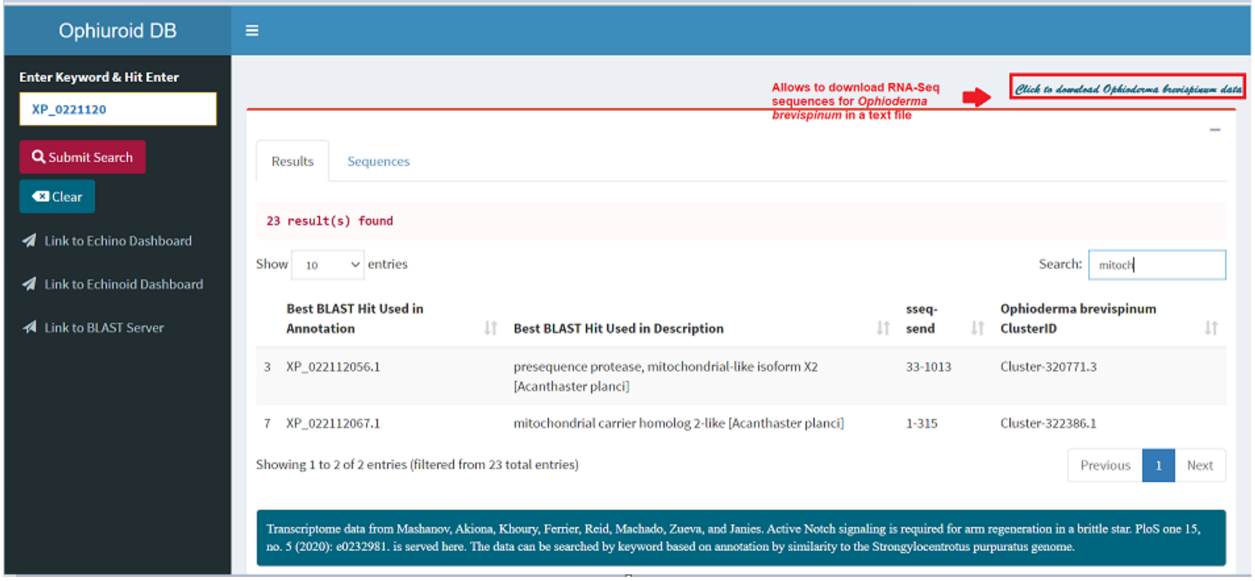


#### 3. Visualizing Sequences

Select a whole row and the record will be highlighted with blue (you can only perform one selection at a time).

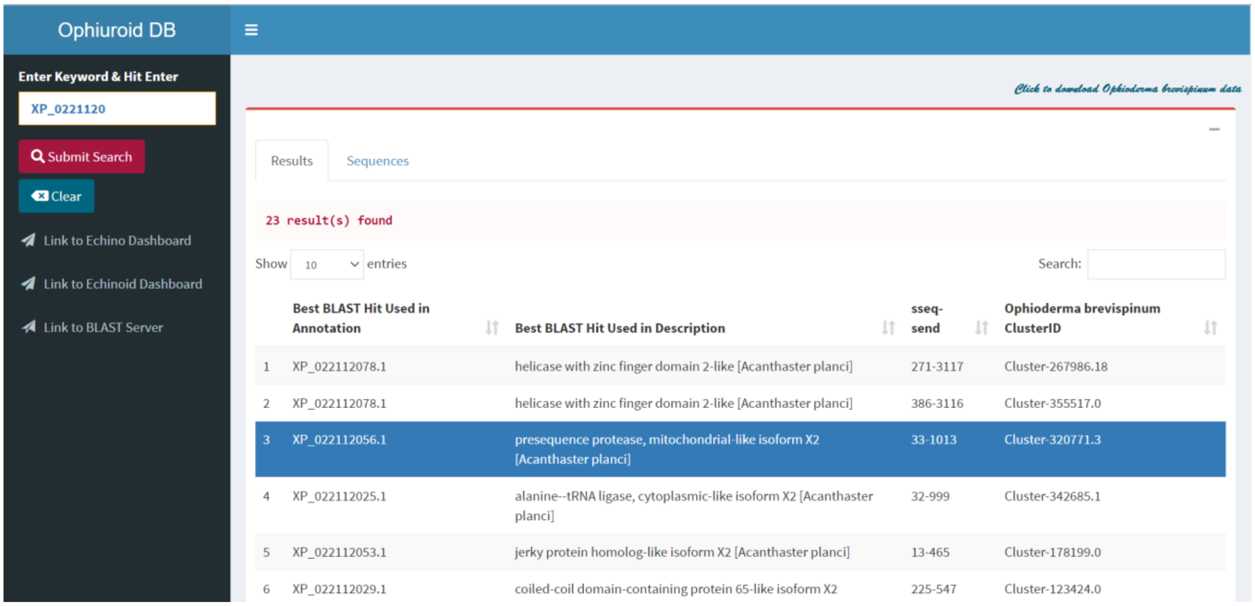

- **Sequences Tab**
  After the record is selected, it redirects you to “Sequences” tab and display protein sequences from Ophiuroids repository. Furthermore, it allows to download result sequence in fasta file format.

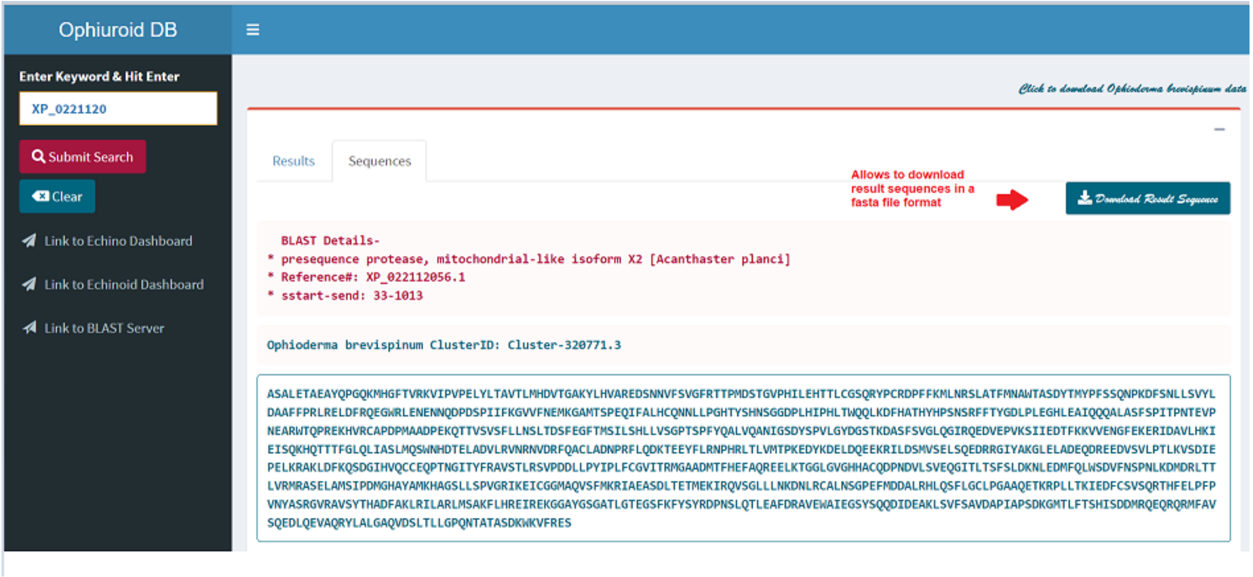


#### 4. Sequenceserver for Basic Local Alignment Search Tool

Access Sequenceserver (Priyam *et al*. 2019) by clicking the “Link to BLAST Server” in the left pane or by type in the following URL: https://echinodb.uncc.edu/sequenceserver/

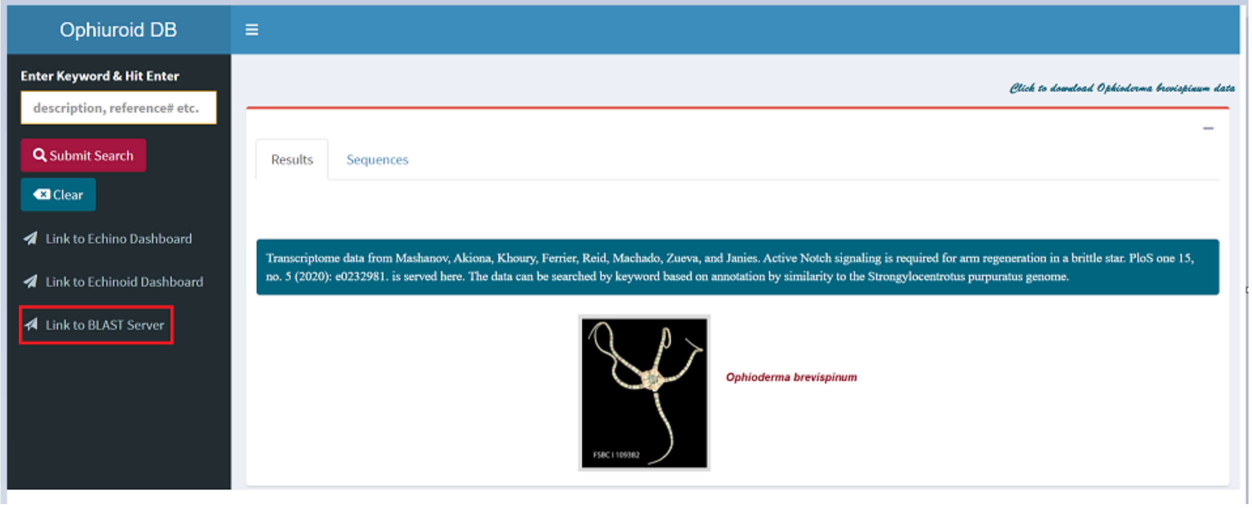

➢ Paste your query string [protein sequence(s)] in the text area to perform BLAST search.

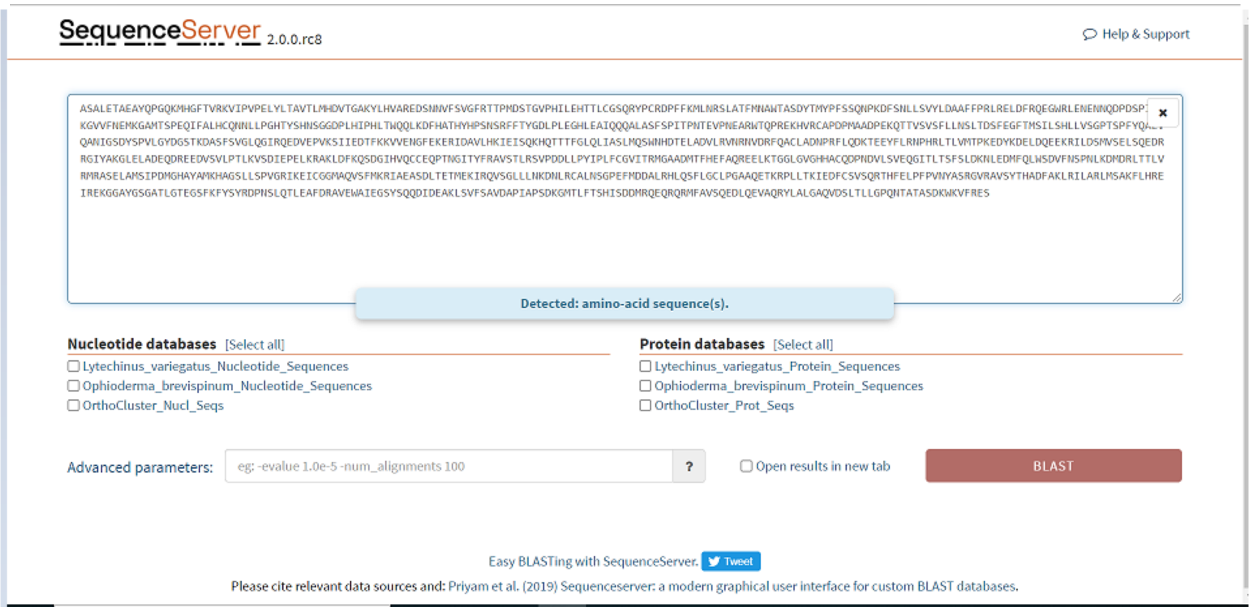


➢ Select database(s) to perform BLAST search against query sequence.

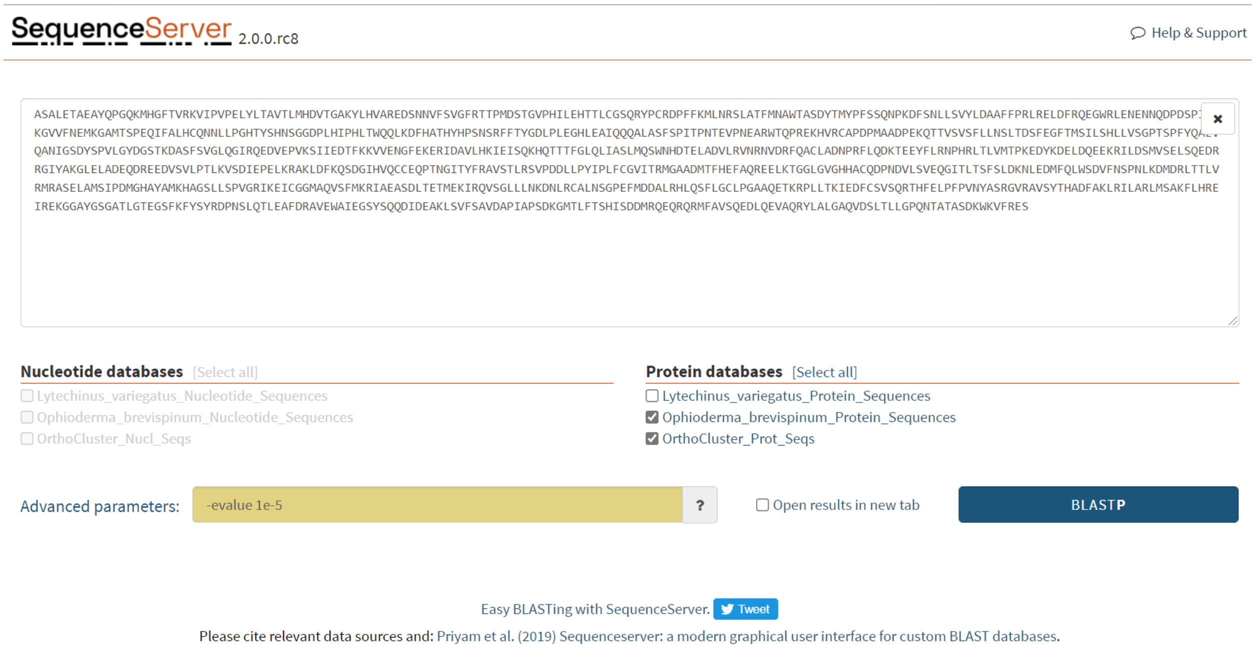

- **Sequenceserver Results**

SequenceServer 2.0.0.rc8

Help & Support

**BLASTP: 1 query, 2 databases**
Edit search | New search

**Download FASTA, XML, TSV**
- FASTA of all hits
- FASTA of selected hit(s)
- Alignment of all hits
- Alignment of selected hit(s)
- Standard tabular report
- Full tabular report
- Full XML report

**SequenceServer 2.0.0.rc8** using **BLAST 2.9.0+**, query submitted on 2021-09-30 01:49:53 UTC
**Databases:** Ophioderma_brevispinum_Protein_Sequences, OrthoCluster_Prot_Seqs (1228855 sequences, 299839407 characters)
**Parameters:** evalue 1e-05, matrix BLOSUM62, gap-open 11, gap-extend 1, filter F
Please cite: https://doi.org/10.1093/molbev/msz185

⊞ **Queries and their top hits: chord diagram**

**Query= Query_1** length: 981

⊟ Graphical overview of hits SVG | PNG

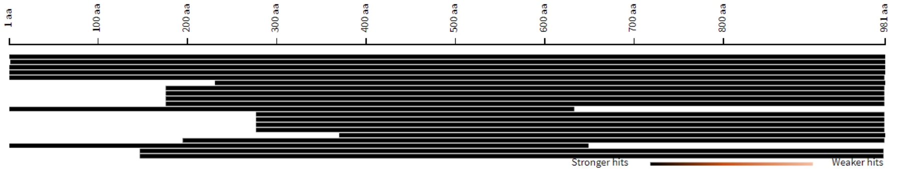


View More

⊞ Length distribution of hits
⊟ Summary table of hits

| # | Similar sequences | Query coverage (%) | Total score | E value | Identity (%) |
|---|---|---|---|---|---|
| 1. | Cluster-320771.3 | 100 | 5313 | 0 | 100 |
| 2. | 20:16798 OS=Psilaster charcoti | 99 | 4616 | 0 | 85 |
| 3. | 19:33421 OS=Peribolaster BJ19 folliculatus | 100 | 4445 | 0 | 82 |
| 4. | 10:37131 OS=Pisaster ochraceus | 100 | 4414 | 0 | 81 |
| 5. | 13:53487 OS=Oligometra serripinna | 99 | 3316 | 0 | 60 |
| 6. | 22:9616 OS=Remaster gourdoni | 77 | 3297 | 0 | 80 |
| 7. | 7:93851 OS=Pteraster tesselatus | 82 | 3244 | 0 | 72 |

- **Download FASTA Sequence, Alignment from a high scoring pair from SequenceServer**

⊟ **20:16798** OS=Psilaster charcoti hit 2, length: 1,003

☐ Select | Sequence | FASTA | Alignment
⊟ Graphical overview of aligning region(s) SVG | PNG

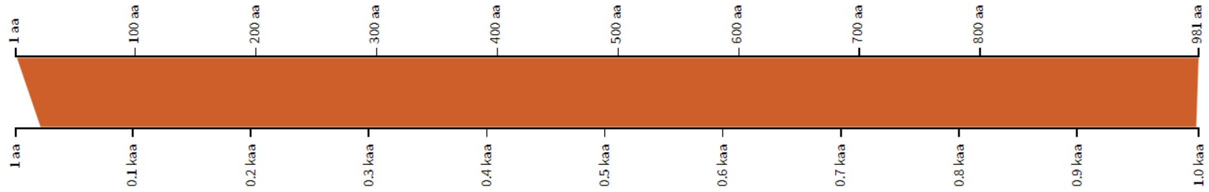


```
Score: 1782.69 (4616), E value: 0, Identity: 833/980 (85%), Positives: 907/980 (92.6%), Gaps: 0/980 (0%)

Query      2 SALETAEAYQPGQKMHGFTVRKVIPVPELYLTAVTLMHDVTGAKYLHVAREDSNNVFSVGFRTTPMDSTGVPHILEHTTLCGSQRYPCRDPFFKMLNR 99
             SAL+TAE Y+PGQK+HGFTV KVIPVPELYLTAV L HDVT AKYLHVAREDSNNVFSV FRTTPMDSTGVPHILEHTTLCGSQRYPCRDPFFKMLNR
Subject   22 SALKTAETYEPGQKLHGFTVEKVIPVPELYLTAVILTHDVTRAKYLHVAREDSNNVFSVSFRTTPMDSTGVPHILEHTTLCGSQRYPCRDPFFKMLNR 119

Query    100 SLATFMNAWTASDYTMYPFSSQNPKDFSNLLSVYLDAAFFPRLRELDFRQEGWRLENENNQDPDSPIIFKGVVFNEMKGAMTSPEQIFALHCQNNLLP 197
             SL+TFMNAWTASDYTMYPFSSQNPKDF NLLSVYLDAAFFPRLRELDFRQEGWRLENENNQDPDSPIIFKGVVFNEMKGAMTSPEQIFA HCQNNLLP
Subject  120 SLSTFMNAWTASDYTMYPFSSQNPKDFKNLLSVYLDAAFFPRLRELDFRQEGWRLENENNQDPDSPIIFKGVVFNEMKGAMTSPEQIFANHCQNNLLP 217

Query    198 GHTYSHNSGGDPLHIPHLTWQQLKDFHATHYHPSNSRFFTYGDLPLEGHLEAIQQQALASFSPITPNTEVPNEARWTQPREKHVRCAPDPMAADPEKQ 295
              HTYSHNSGGDPLHIP LTWQQLK+FHATHYHPSNSRFFTYGDLPLEGHLEAIQQQAL SFSPI+P+TEVPNE RWTQPREKHV CAPDPMAADPEKQ
Subject  218 SHTYSHNSGGDPLHIPRLTWQQLKEFHATHYHPSNSRFFTYGDLPLEGHLEAIQQQALESFSPISPDTEVPNETRWTQPREKHVHCAPDPMAADPEKQ 315

Query    296 TTVSVSFLLNSLTDSFEGFTMSILSHLLVSGPTSPFYQALVQANIGSDYSPVLGYDGSTKDASFSVGLQGIRQEDVEPVKSIIEDTFKKVVENGFEKE 393
             TTVSVSFLLNSL DSFEGFT+SILSHLLVSGPTSPFYQAL++ANIGSDYSPVLGYDGSTKDASFSVGLQGIR+EDVE VK II DTFKKVVE GFEKE
Subject  316 TTVSVSFLLNSLIDSFEGFTLSILSHLLVSGPTSPFYQALLEANIGSDYSPVLGYDGSTKDASFSVGLQGIREEDVEKVKGIIHDTFKKVVETGFEKE 413

Query    394 RIDAVLHKIEISQKHQTTTFGLQLIASLMQSWNHDTELADVLRVNRNVDRFQACLADNPRFLQDKTEEYFLRNPHRLTLVMTPKEDYKDELDQEEKRI 491
             RIDA+LHKIEISQKHQ+TTFGL L+ASLMQSWNHD +LA ++RVN+NV+RFQACLADNPRFLQDKTEEYF  N HRLTLVMTPKEDY + +DQEEKRI
Subject  414 RIDAILHKIEISQKHQSTTFGLNLVASLMQSWNHDADLASIMRVNKNVERFQACLADNPRFLQDKTEEYFTSNNHRLTLVMTPKEDYNETIDQEEKRI 511
```

#### 5. Clear Search/Results in Ophiuroid DB

Go to the results tab and hit “Clear” button or “Delete” key to clear the search.

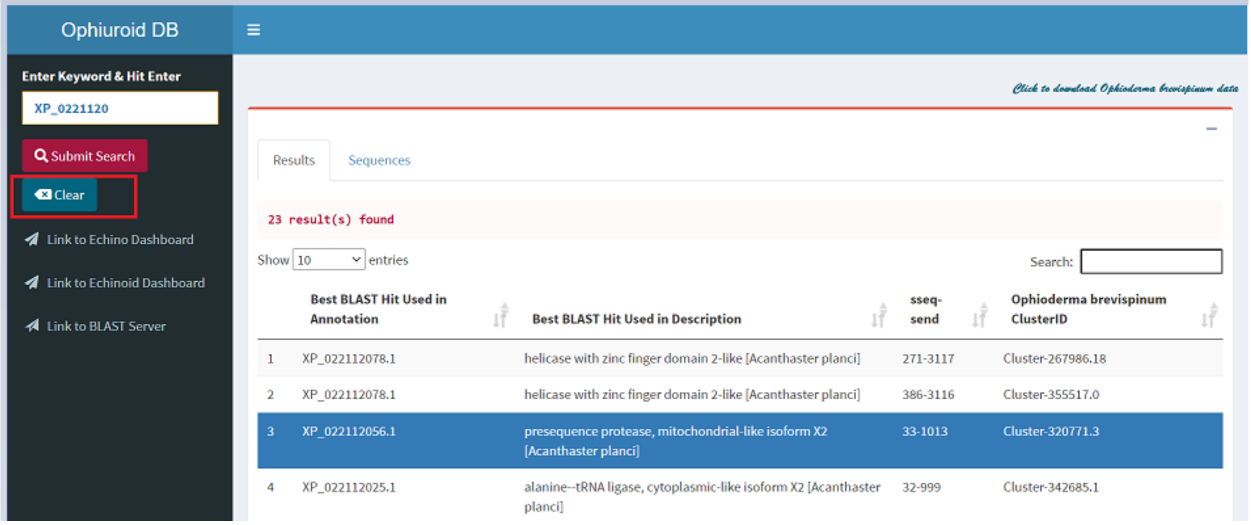


- And, search will be cleared after the button is clicked or delete key is pressed.

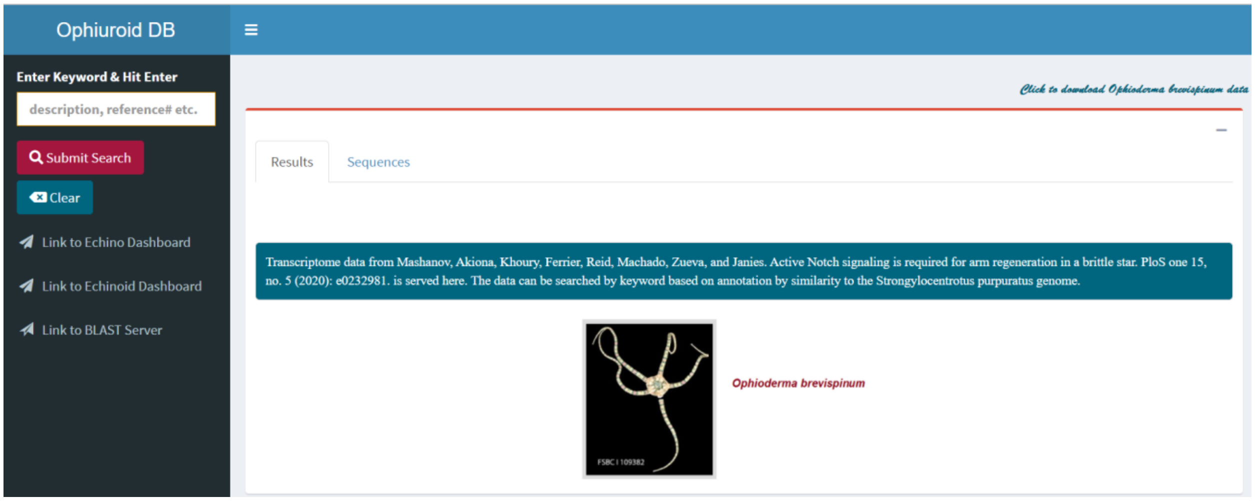

#### 6. Additional Links

Links in the left pane are provided to redirect users to “EchinoDB” or “EchinoidDB” page.

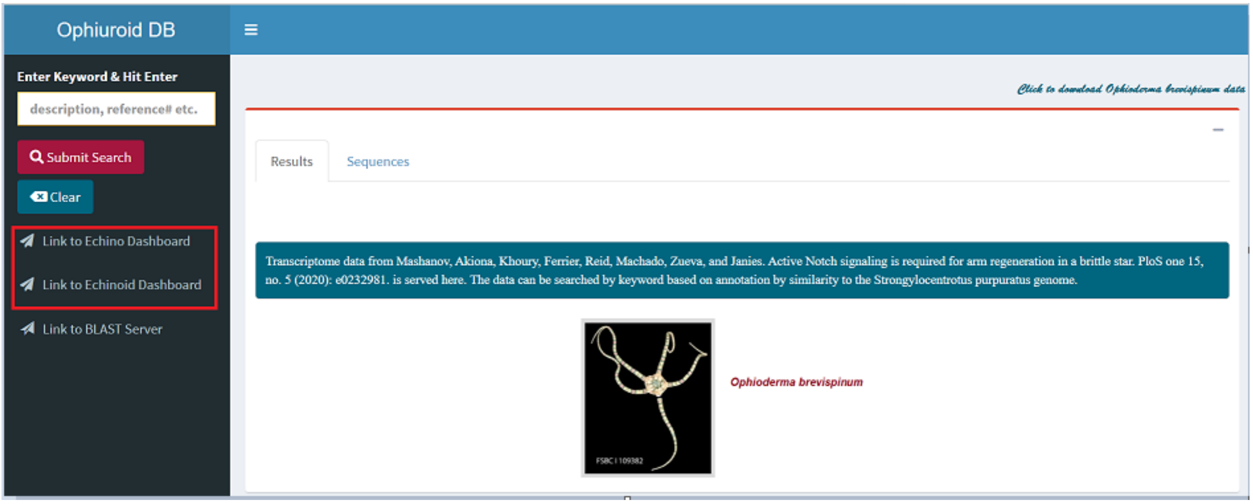
